# Functional characterization of RNA profiles in processing bodies of human embryonic stem cells and mesodermal cells

**DOI:** 10.1101/2023.11.22.568232

**Authors:** Jin Jiang, Qizhe Shao, Sisi Xie, Xiaoying Xiao, Ruisi Guo, Min Jin, Di Chen

**Author notes:** Corresponding author. (DC) (JJ).

## Abstract

Processing bodies (P-bodies) are the membraneless organelles that play critical roles in RNA storage and decay. Abnormalities in P-bodies contribute to diseases and developmental disorders. Huge efforts have been applied to the identification of the protein components in P-bodies, however, the dynamics of RNA components of P-bodies in human embryonic stem cells (hESCs) during maintenance and differentiation remain largely elusive. Here, we characterized the RNA profiles of P-bodies from HEK293T cells, hESCs, and hESC-derived mesodermal cells. The number of P-bodies decreases upon hESC differentiation towards mesodermal fate, accompanied with the decreased RNAs within P-bodies. By functional analysis of the P-body enriched and P-body depleted genes across different cell types, we discovered the cell type-specific enrichment of P-body-genes and the potential association with human diseases. We also captured the non-coding RNAs, including long intergenic non-coding RNAs and transposable elements in P-bodies in a cell type-specific manner. Furthermore, we verified the involvement P-bodies in regulating the differentiation of hESCs towards mesoderm by over-expression of LSM14A and knocking down of *SPTAN1*. In summary, we characterized the mRNAs and non-coding RNAs in P-bodies of hESCs and hESC-derived mesodermal cells, discovering the potential roles that P-bodies play for the precise differentiation of hESCs.

## INTRODUCTION

Pluripotent stem cells (PSCs), including embryonic stem cells (ESCs) and induced pluripotent stem cells (iPSCs), possess the capacity to self-renew to maintenance themselves as well as the capability to differentiate into different lineages of functional cells triggered by different signals^1,2^. The differentiation of PSCs is precisely regulated by different mechanisms to achieve meticulous gene expression programs for specific cell fates, including regulation at transcriptional, post-transcriptional, translational, and post-translational levels^3,4^. Transcriptional regulation is mainly based on the interaction of the local epigenetic landscape and transcription factors/regulators that either promote or repress transcription machinery for gene expression. Key transcription factors, such as OCT4, KLF4, and NANOG, have been identified to play key roles to function cooperatively in regulating the maintenance and differentiation of PSCs^5,6^. Notably, these transcription factors participate in the organization of chromatin via liquid-liquid phase separation (LLPS) to regulate the expression of pluripotency and differentiation-related genes^7,8^.

Regulation via forming condensates not only happens in the nucleus for transcription, but also exists in cytoplasm for post-transcriptional modulation. Recent studies of intracellular phase separation have extended our understanding of biomolecular condensates involved in the orchestration of cell fates^9,10^. Like OCT4 condensates, RNA-binding proteins (RBPs) could also form a functional hub through phase separation by recruiting RNAs and other RBPs. These RBP-centered condensates play key roles in post-transcriptional regulation of RNAs, such as splicing, alternative polyadenylation, cellular localization, and stability^9,11^. For example, FXR1 recruits translation machinery to activate translation of the stored mRNA in FXR1 granules formed by LLPS in spermatid stage to ensure spermatogenesis ^12^. Besides, there are different kinds of granules or condensates formed by LLPS of RBPs and RNAs that regulate the RNA metabolism for coordinating cell fates in physiological and/or different stressed conditions, such as processing bodies (P-bodies) and stressed granules^9,13–15^.

P-bodies are cytoplasmic membraneless organelles formed by LLPS that play critical roles in regulating RNAs at post-transcriptional level^10,16^. Conserved from yeast to mammals, P-bodies are the sites for mRNA storage and/or decay, mainly related to the repression of mRNA translation^16–20^. Importantly, mRNAs stored in P-bodies could be translocated to ribosomes for translation, indicating that mRNAs are not permanently silenced or decay in P-bodies, but rather are highly dynamic and regulated^21^. Recent studies have demonstrated that DDX6 (Patr-1 in Drosophila), the key components of P-bodies, is critical for the maintenance of pluripotent states of human PSCs and intestinal stem cells in Drosophila^22,23^. Moreover, dysregulation of P-bodies have been discovered to contribute to neural diseases, cancers, and developmental disorders^9,24,25^. However, the dynamics of P-body components during development remain largely unknown and the functions of P-body-mediated regulation in development are still elusive.

In this study, we applied fluorescence-activated particle sorting to purified P-bodies from 293T, human ESCs (hESCs), and mesodermal cells induced from hESCs. We analyzed the RNA components of P-bodies in different cell types and discovered the common and unique features of P-bodies that may play critical roles for the regulation of stem cell fate and differentiation.

## RESULTS

### Purification of processing bodies from HEK293T cells, human embryonic stem cells, and mesodermal cells

Processing bodies (P-bodies) are one of the membraneless organelles in cytoplasm that play critical roles in RNA regulation^10,16,26^. Characterization of P-body components is key to understanding the functions of P-bodies, especially during cell fate transition. To systematically investigate the potential functions of P-bodies in regulating the maintenance and differentiation of human embryonic stem cells (hESCs), we applied the fluorescent associate particle sorting (FAPS) strategy^19^ to isolate the P-bodies from hESCs and hESC-derived mesodermal cells (meso). We also included HEK293T (293T) cells for comparison (Figure 1A). We aimed to identify hESC-specific components and mesoderm-specific components of P-bodies and uncover the potential functions of P-bodies during the maintenance and differentiation of hESCs.

**Figure 1.**
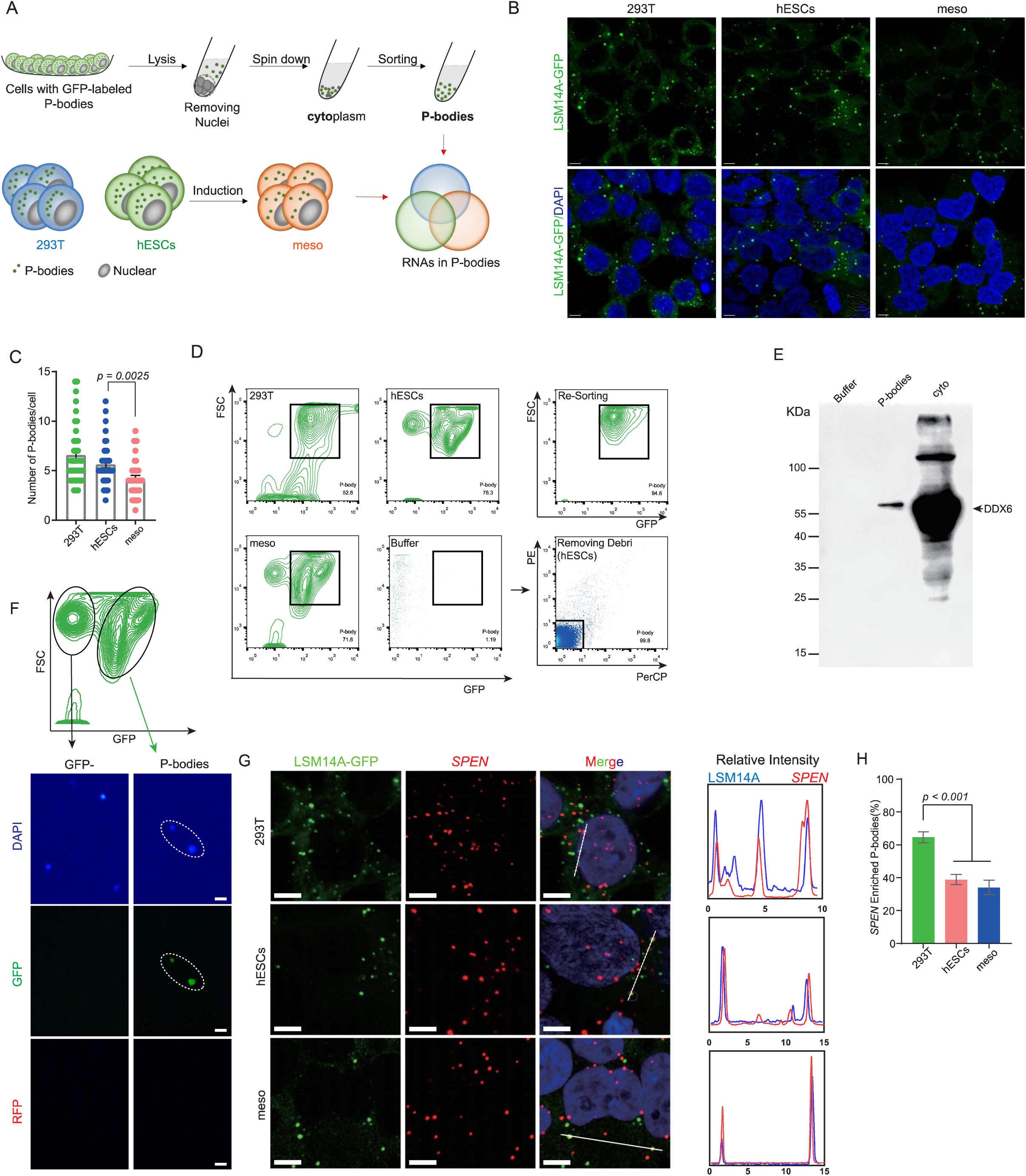
Purification of P-bodies from HEK293T cells, hESCs, and hESC-derived mesodermal cells. (A) Schematic illustration of the experimental design. P-bodies are purified by fluorescence-activated particle sorting from HEK293T cells (293T), hESCs and hESC-derived mesodermal cells (meso) for analysis. (B) Confocal images showing the LSM14A-GFP-labeled P-bodies in 293T, hESCs, and meso. Scale bar: 5 μm. (C) Quantification of P-bodies in 293T, hESCs, and meso. N = 40 cells for each group; error bars represent mean ± SEM. The numbers of P-bodies per cell are: 6.59 ± 0.48 for 293T; 5.64 ± 0.35 for hESCs; 4.20 ± 0.30 for meso. (D) Fluorescence-activated particle sorting of P-bodies from 293T, hESCs, and meso based on their size (FSC) and GFP fluorescence intensity. PE and PerCP channels are used to exclude cellular debris. (E) Western blot analysis of buffer control, purified P-bodies, and cytoplasm (cyto) using anti-DDX6 antibody. (F) Epifluorescence imaging of sorted GFP positive P-bodies and GFP negative particles. LSM14A-GFP labeled P-bodies are highlighted by the white dashed circle. DAPI staining (blue) is applied to indicate nucleic acids. RFP channel is used to exclude the influence of auto-fluorescence. Scale bar: 1 μm. (G) RNAscope images showing the enrichment of *SPEN* mRNAs (red) in P-bodies labeled with LSM14A-GFP in 293T, hESCs and meso. DAPI (blue) is countered stained to indicate nuclei. Line scans are employed to show the related intensity profiles of LSM14A-GFP and *SPEN* mRNAs on the right panel. scale bar: 5μm. (H) The percentage of P-bodies containing *SPEN* mRNAs (N = 40 cells, data is shown in mean±SEM. 293T: 64.56±3.318%; hESCs: 38.85±3.106%; meso: 34.07±4.396%). **See also Figure S1 and S2.**

To establish a stable and reliable fluorescent marker for sorting P-bodies, we first tested the commonly used P-body markers including LSM14A, DDX6, and DCP1A, by using an inducible overexpression system^10,27^. Fluorescence-labeled P-bodies could be visualized after adding doxycycline (Dox) for 4 to 6 days (Figure S1A). Among these three reporters, LSM14A-GFP resembles P-bodies in morphology and number (Figure S1A). To test the persistency of the three P-body reporters, the GFP-positive cells were sorted into three groups, high, intermediate, and low, based on GFP intensity. Notably, P-bodies were only found in the high GFP intensity cells. After three days of culture, LSM14A showed high proportions of GFP-positive P-bodies, whereas DDX6 and DCP1A labeled cells showed declined GFP-positive P-bodies (Figure S1B). Next, we determined the theoretical isoelectric points for LSM14A, DDX6, and DCP1A by Expasy prediction (https://www.expasy.org/), and discovered that LSM14A is the highest (Figure S1C), suggesting that LSM14A is probably the best candidate for P-body formation^11,28^. This is supported by the prediction of intrinsically disordered region (IDR) by PONDR (http://www.pondr.com/) and Alphafold2 (https://alphafold.com/) (Figures S1C-D). Therefore, LSM14A-GFP represents the best reporter for labeling P-bodies. To trace the dynamics of P-bodies using LSM14A-GFP, we added the P-body inhibitor cycloheximide (CHX)^17,29^ and observed the disappearance of GFP-labeled P-bodies. Importantly, GFP-labeled P-bodies re-appeared after washing out CHX (Figure S1E). This is also confirmed by Fluorescence recovery after photobleaching (FRAP) (Figure S1F)^18^. Based on all the observations above, we chose LSM14A-GFP as the P-body reporter for subsequent experiments and analyses.

Next, we verified the LSM14A-GFP reporter expression in hESCs by immunofluorescence for P-body markers DDX6 and DCP1A (Figure S2A). Overexpression of LSM14A did not change the expression of other P-body markers as expected (Figure S2B). We chose mesoderm induction to represent hESC differentiation for exploring the P-body dynamics. Importantly, over-expression of LSM14A had no obvious impact on the expression of the pluripotency or differentiation-related genes (Figure S2C-F), therefore, we could use this LSM14A-GFP to explore the P-body dynamics in hESC maintenance and differentiation.

Based on these discoveries, we made LSM14A-GFP reporters in 293T cells, hESCs, and hESC-derived mesodermal cells (meso) (Figure 1B). The number of P-bodies per cell in 293T was consistent with previous report^30^. Interestingly, the number of P-bodies per cell in hESCs is comparable to that of 293T cells, but slightly reduced in meso (Figure 1B-C). We then purified P-bodies from 293T cells, hESCs, and meso with LSM14A-GFP (Figures 1A, D). We verified the purified P-bodies by western blot examination of P-body marker DDX6^10,27^ (Figure 1E) and DAPI staining (Figure 1F). In addition, we detected the presence of *SPEN* mRNA, a known P-body component^19^, in P-bodies using RNAscope. *SPEN* mRNAs were present in the P-bodies in 293T cells, however, the proportion of *SPEN* in the P-bodies of hESCs and meso was relatively lower compared to 293T cells (Figures 1G-H, Figure S2G-H), indicating the potential divergence of P-body components in different cell types. Taken together, LSM14A-GFP was identified as the most suitable marker for sorting P-bodies from 293T, hESCs and meso.

### Characterization of RNA components in P-bodies from different cell types

To characterize the RNA components in P-bodies across different cell types, we performed low-input RNA-sequencing on the sorted P-bodies and the corresponding cytoplasm (cyto) fractions from 293T, hESCs, and meso. The replicates from each sample are highly correlated (Figure 2A and Figure S3A). Different fractions from the same cell type are clustered together based on Principle Components Analysis and correlation analysis (Figures S3B-C). Notably, P-bodies from different cell types are clustered together based on hierarchical clustering analysis (Figure 2B), indicating that the RNAs in the P-bodies may share significant similarity among these three cell types. Next, we determined the P-body enriched genes and P-body depleted genes by analyzing the differentially expressed genes (DEGs) in P-body fraction and cyto fraction. We identified 918 P-body enriched genes and 991 P-body depleted genes in 293T cells, 1456 P-body enriched genes and 971 P-body depleted genes in hESCs, and 569 P-body enriched genes and 518 P-body depleted genes in meso (Figures 2C, E, Table S1). We also determined the P-body enriched and P-body depleted long non-coding RNAs (lncRNAs) and discovered the similar trends as mRNAs (Figure 2D and Figure S3D, E, Table S2).

**Figure 2.**
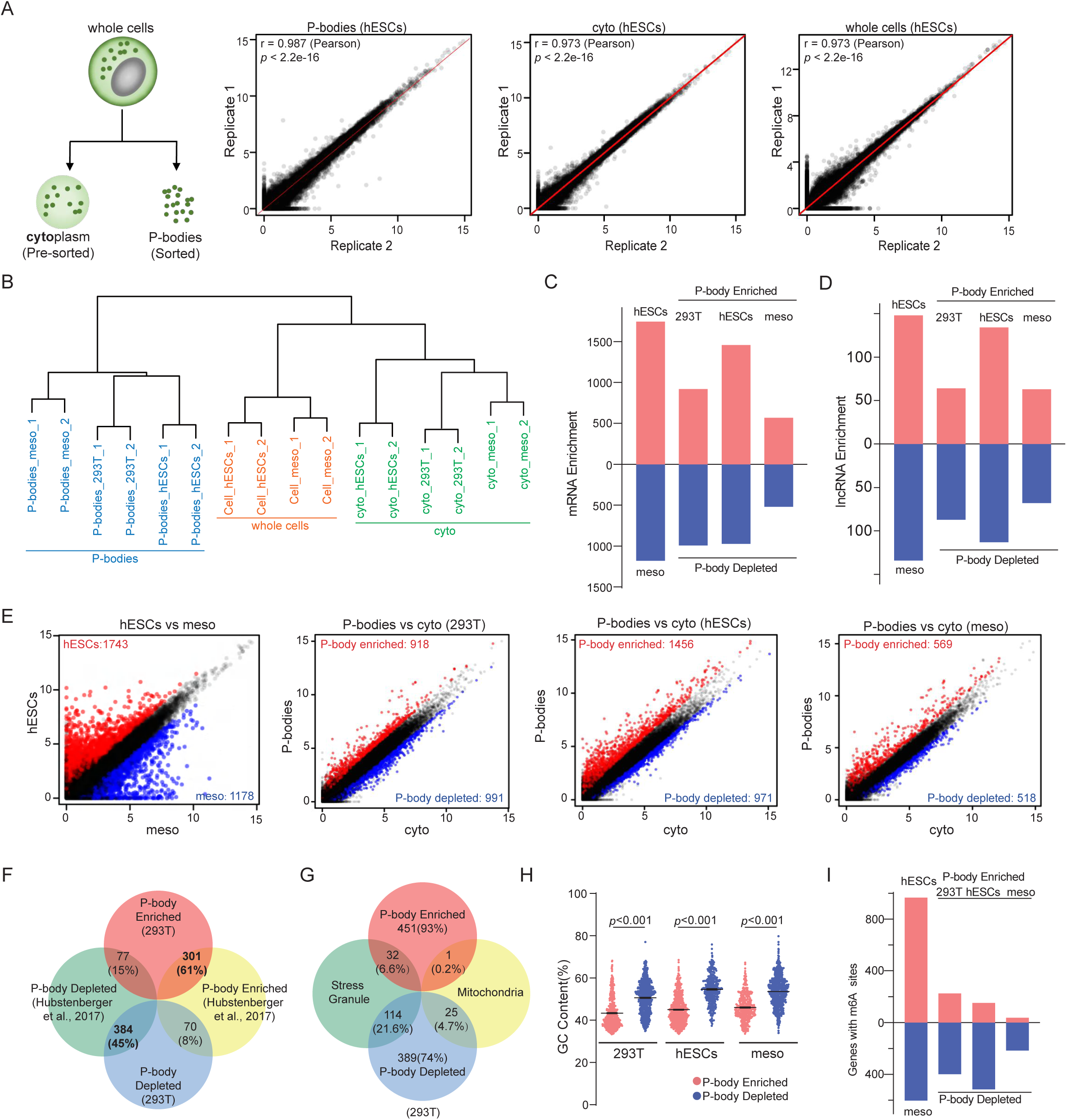
Characterization of the RNAs in P-bodies. (A) Scatter plots showing the biological replicates of the transcriptomes from purified P-bodies, cyto (the pre-sorted fractions), and the whole cells of hESCs. Pearson test was conducted. (B) Unsupervised hierarchical clustering analysis of the transcriptomes of the whole cells, cyto, and P-bodies from 293T, hESCs, and meso. (C-D) Bar plots showing the enriched genes for hESCs vs meso, and P-body enriched genes and depleted genes in 293T, hESCs, and meso of mRNAs (C) and lncRNAs (D). (E) Scatter plots showing the comparisons of the DEGs in hESCs vs meso, P-body vs cyto of 293T, P-body vs cyto of hESCs, and P-body vs cyto of meso. (F) Venn plots showing the comparisons of P-body enriched and depleted genes in 293T cells from this study, P-body enriched genes and P-body depleted genes from Hubstenberger et al., 2017^19^. (G) Venn plots showing the comparisons of P-body enriched genes and P-body depleted genes, to genes in stress granules^50^ and genes in mitochondria^51,52^ in 293T cells. (H) The distribution of the GC contents of the P-body enriched (red) and P-body depleted (blue) genes in 293T, hESCs, and meso (Data is shown in mean±SEM. 293T: 43.4±0.26 for P-body enriched group and 53.6±0.23 for P-body depleted group. hESCs: 45±0.2 for P-body enriched group and 50.6±0.2 for P-body depleted group. Meso: 46±0.28 for P-body enriched group and 54.6±0.32 for P-body depleted group). (I) Bar plots showing the number of mRNAs with m^6^A modification sites for hESC-enriched and meso-enriched genes, and for P-body enriched and P-body depleted genes of 293T, hESCs, and meso. **See also Figure S3.**

We next compared the P-body enriched genes and P-body depleted genes in 293T cells from our study to previously published data^19^. Importantly, 61% of the P-body enriched genes and 45% of the P-body depleted genes from this study were also detected in the published study^19^, but not overlapping with genes in mitochondria or stress granules (Figures 2F-G), confirming our purified P-bodies are suitable for deep analysis. Furthermore, the GC contents of the mRNAs are lower compared with mRNAs in cyto of 293T cells, consistent with previous finding ^20^, and the same trend for hESCs and meso (Figure 2H and Figures S3G-I). Given that *N*6-methyladnisine (m^6^A) is critical for regulating mRNA fate^31,32^, we also examined the m^6^A modification in P-body enriched and P-body depleted genes. Interestingly, genes with m^6^A modification are enriched in P-body depleted fraction, and the m^6^A sites per gene were slightly lower in P-body enriched genes (Figure 2I, Figure S3J). Taken together, we characterized the RNA profiles in P-bodies from 293T cells, hESCs, and meso, forming the basis for analyzing P-body enriched and P-body depleted genes related to cellular functions.

### Cell type-specific enrichment of mRNAs and lncRNAs in P-bodies

To further analyze the potential functions of the RNAs in P-bodies across different cell types, we compared the P-body enriched genes in 293T, hESCs, and meso (Figure 3A).

**Figure 3.**
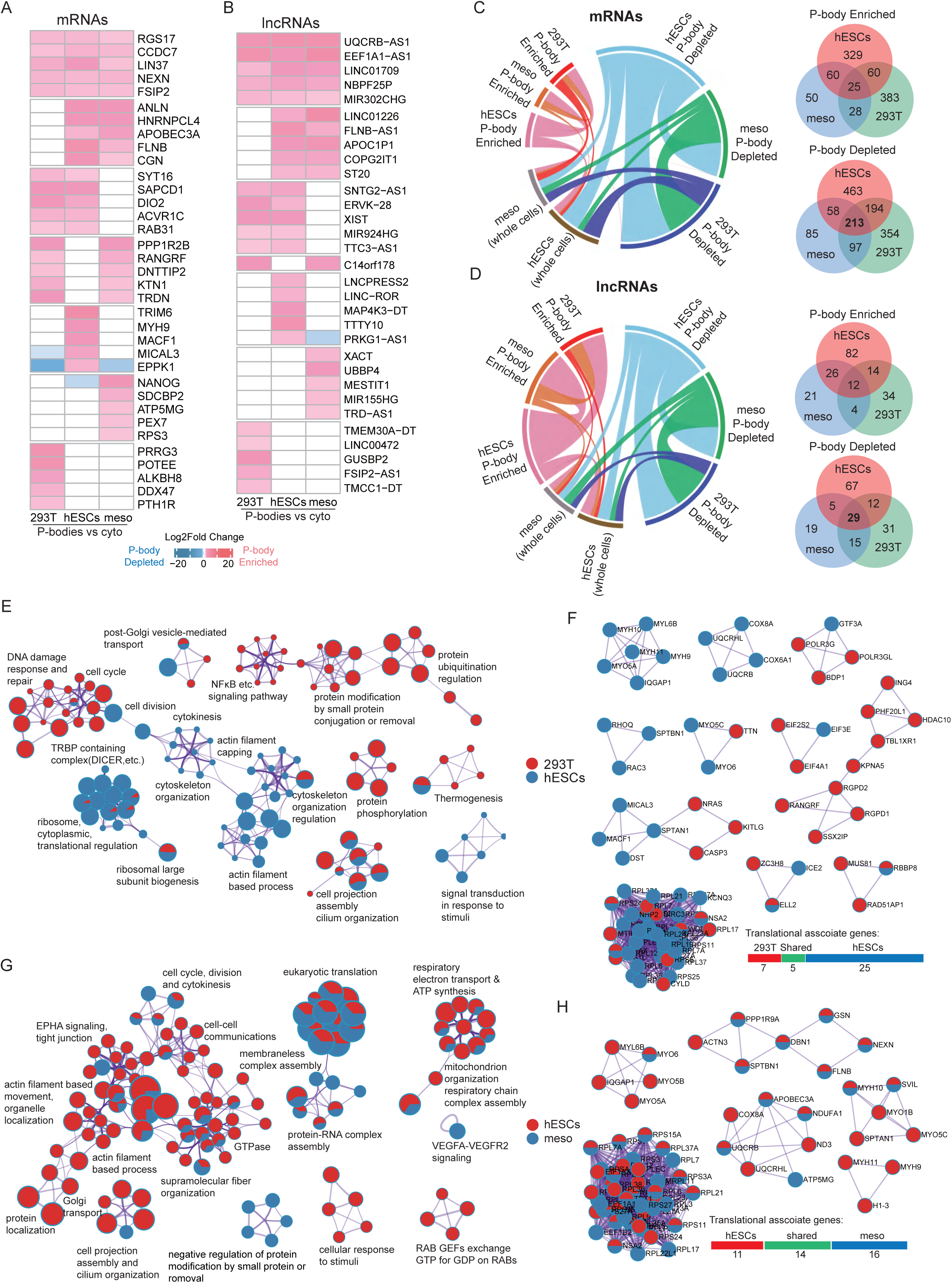
The properties of P-body enriched and P-body depleted genes in 293T cells, hESCs, and hESC-derived mesodermal cells. (A) Heatmap showing the P-body enriched mRNAs shared or unique for different cell type(s) of 293T, hESCs, and meso. (B) Heatmap showing the P-body enriched lncRNAs shared or unique for different cell type(s) of 293T, hESCs, and meso. (C-D) Circos (left) and Venn (right) plots showing the overlaps of P-body enriched and P-body depleted mRNAs (C) and lncRNAs (D). (E) Functionally enriched clusters showing the P-body properties for 293T (red) and hESCs (blue). (F) Core protein-protein interaction (PPI) networks showing the potential regulatory sets mediated by P-bodies in 293T (red) and hESCs (blue). (G) Functionally enriched clusters showing the P-body properties for hESCs (red) and meso (blue). (H) Core PPI network showing the potential regulatory sets mediated by P-bodies in hESCs (read) and meso (blue). **See also Figure S4 and S5**

We identified P-body enriched mRNAs encoding cytoskeleton (NEXN, ANLN, CGN, MYH9, FLNB), GTPase (RGS17, RAB31), protein modifications (DIO2, APOBEC3A, TRIM6), and RNA binding proteins (SYT16, HNRNPCL4) that are shared or unique by different cell types (Figure 3A). This is consistent with previous findings that P-bodies play key roles in RNA metabolism and protein modifications and regulation^10,16,19^. Interestingly, the pluripotency gene NANOG is enriched in the P-bodies of meso, but in contrast, depleted in P-bodies of hESCs, suggesting that P-bodies may be involved in the clearance of NANOG in meso (Figure 3A). Similarly, lncRNAs also exhibit cell type-shared and -specific manner in P-bodies of 293T, hESCs, and meso (Figure 3B). Furthermore, there was a remarkable overlap of P-body depleted RNAs, including mRNAs and lncRNAs, among the three cell types (Figures 3C-D), suggesting that P-body enriched RNAs are cell-type specific, while the RNAs in the cyto (P-body depleted) are relatively similar in different cell types.

To further elucidate the potential cell-type-specific enrichment of P-body mRNAs, we first compared the P-body enriched mRNAs in 293T and hESCs. Importantly, 293T cells and hESCs shared terms such as protein localization, cell projection, and RNA-protein complex formation including membraneless organelles and Golgi apparatus transport functions. Interestingly, hESCs enriched terms such as actin organization or function, translational associate or ribosome formation, cytoskeleton, stimuli response, and cytokinesis functions. While 293T cells enriched terms including protein phosphorylation, modification, ubiquitination regulation, and DNA damage repair response functions (Figure 3E and Figures S4A-B). We then analyzed the protein-protein interactions (PPIs) of the P-body enriched genes in 293T cells and hESCs. Interestingly, 293T and hESC P-body enriched genes are interacting tightly within each cell type, further reflecting the cell-type-specific functions of P-bodies (Figure 3F and Figure S4C).

Next, we compared the P-body enriched mRNAs in hESCs and meso to uncover the potential regulatory functions of P-bodies during hESC differentiation. hESCs and meso shared terms such as translation regulation, RNA-protein complex formation including membraneless organelles, cytoskeleton organization and functions, cytokinesis, cell cycle and division, GTPase, ATP synthesis, cell projection associate genes. hESCs enriched terms including stimuli response, action filament process and movement, organelle localization, protein localization, Golgi mediate transport, stimuli response, cell-cell communications associated functions. While meso enriched protein modification related terms (Figure 3G and Figure S4D), these functions are mainly related to metabolic and cellular processes for both hESCs and mesodermal cells (Figure S4E). PPI analysis revealed that a significant proportion of the interacting genes are shared by hESCs and meso, or specific to hESCs, and most of the PPIs within meso enriched genes are related to protein translational regulation (Figure 3H, Figure S4F). Notably, nearly 40% of the P-body enriched genes of meso are related with translational regulation. Furthermore, translational regulation related genes are the most abundant, although the expression level of these genes is relatively low (Figure 3H, Figures S4G-H). These observations suggest that the transition from hESCs to meso involves specific changes in the composition and function of P-bodies, particularly with respect to translation-related processes.

We then analyzed the P-body depleted genes in different cell types (Figures S5A-B). Some embryonic development-related genes, such as MIXL1, GATA3, SALL3, WNT6, BMP3, and FOXO6, are depleted in P-bodies of shared or specific cell types. Interestingly, the P-body depleted genes from different cell types were almost enriched in identical or similar terms, mainly related to development, metabolism and cellular processes (Figures S5C-E).

### Functional analysis of P-bodies in different cell types

The cellular functions of proteins are highly determined by their subcellular localization^33^. Therefore, we examined the subcellular localization of the proteins encoded by the cell-type-specific P-body enriched mRNAs. Although more DEGs are found in P-bodies of hESC (Figure 2C), less terms were detected compared to 293T or meso (Figure 4A). To investigate the possible roles of P-bodies in stem cell differentiation, we focused on the GO terms and KEGG pathways related to stem cell development. Interestingly, although the P-body enriched genes did not show significant associations with these processes, the P-body depleted genes were highly associated with stem cell development (Figure 4B), which is consistent with the finding that P-body depleted genes are highly related to development (Figure S5C). These observations promoted us to explore whether P-body enriched or P-body depleted genes are associated with human diseases. Consistently, P-body depleted genes are relatively more associated with human diseases such as cancers, developmental disorders, and neurodegeneration-related diseases in all three cell types (Figures 4C-E and Table S3). Furthermore, these disease-associated genes enriched or depleted in P-bodies are related with similar GO terms to the P-body enriched or P-body depleted genes as mentioned above (Figure 4F). Taken together, these results suggest that cellular critical genes are more excluded from P-bodies, while P-bodies enrich more cell-type-specific genes for different cell fates.

**Figure 4.**
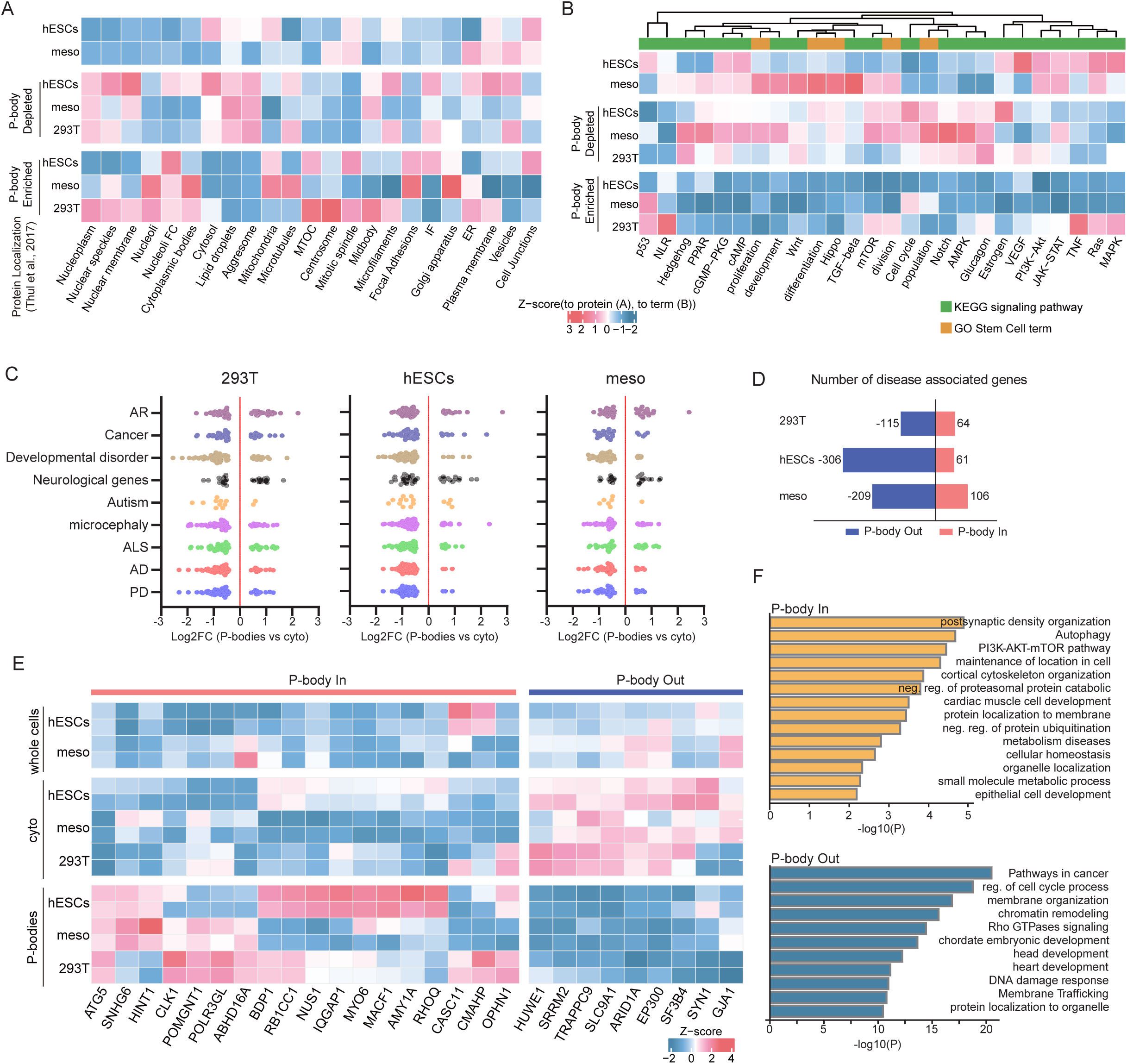
Functional analysis of mRNAs enriched or depleted in P-bodies. (A) Heatmap showing the sub-cellular localization of the proteins encoded by the P-body enriched or P-body depleted genes in 293T, hESCs, and meso. (B) Heatmap showing the enrichments of the key signaling pathways of KEGG terms and stem cell-related GO terms for P-body enriched and P-body depleted genes in 293T, hESCs, and meso. NLR: NOD-like receptors. PPAR: peroxisome proliferator activated receptors. AMPK: Adenosine 5’-monophosphate (AMP)-activated protein kinase. VEGF: vascular endothelial growth factors. TNF: tumor necrosis factor. MAPK: mitogen-activated protein kinase. (C) Dot plot showing the expression of the diseases-associated genes in P-bodies (Fold Change >0, FDR <0.05) or in cyto (Fold Change <0, FDR <0.05) of 293T, hESCs and meso (FDR <0.05). Disease-associate gene sets are derived from OMIM database (https://www.omim.org/). AR: autosomal recessive inheritance. ALS: amyotrophic lateral sclerosis. AD: Alzheimer’s disease. PD: Parkinson’s disease. (D) Bar plot showing the numbers of disease-associated genes in P-bodies (log2FC >0, FDR <0.05) or out of P-bodies (log2FC <0, FDR <0.05). (E) Heatmap showing the enrichment of the disease-associated genes in P-bodies, cyto, or the whole cells of 293T, hESCs or meso. (F) Bar plots showing the functional enrichment of disease-associated genes in (top, orange) or out of (bottom, blue) P-bodies.

### Analysis of non-coding RNAs in P-bodies from different cell types

Next, we analyzed the non-coding RNAs in P-bodies. LncRNAs are involved in chromatin remodeling, transcriptional regulation, and mRNA metabolism by interacting with DNA, RNA, or protein^34^. Therefore, we first analyzed the protein and RNA interactions of P-body enriched or depleted lncRNAs. We discovered that P-body depleted lncRNAs showed a tendency to interact with proteins, while P-body enriched lncRNAs showed more interactions with RNA (Figures 5A, B). Protein-lncRNA interactions were comparable in P-bodies and cytoplasm of hESCs, in contrast to 293T cells or meso where protein-lncRNA interactions are more excluded from P-bodies (Figures 5A, B). Transcription factor (TF) networks play key roles in the regulation of pluripotency^35^. Based on the TF perturbation analysis, we found that TFs associated with lncRNA regulation are highly enriched in P-bodies (Figure 5C), further supporting the finding that RNAs are under tight regulation. Notably, these lncRNA-associated TFs include the pluripotency regulator NANOG, SOX2, and OCT4 (Figure S6). Taken together, these observations highlight the divergence of lncRNAs in P-bodies and cytoplasm, implying the potential lncRNA-mediated post-transcriptional regulation in P-bodies that play important roles in the regulation of stem cells.

**Figure 5.**
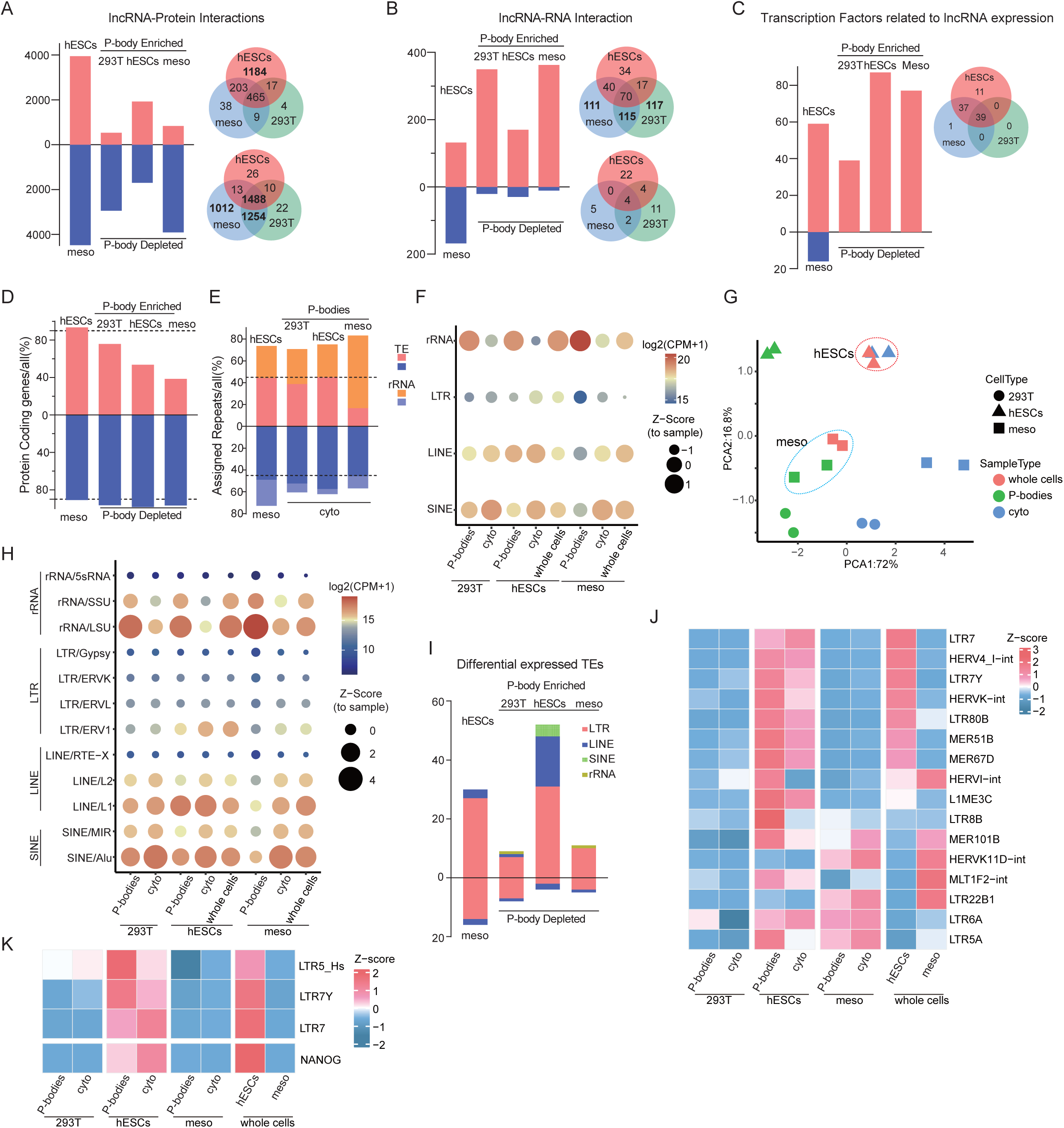
Analysis of lncRNAs, rRNAs, and transposable elements in P-bodies. (A-C) Bar plots showing the lncRNAs interacting networks including: protein-lncRNA pairs (A), RNA-lncRNA pairs (B), and the number of transcription factors related to the expression of lncRNA based on lncSEA (https://bio.liclab.net/LncSEA/index.php) (D) Bar plot showing the percentages of P-body enriched or P-body depleted protein coding genes relative to all genes from 293T, hESCs, and meso. Gene types were determined by ensembl bioMart tool. (E) Bar plot showing the percentages of P-body enriched or P-body depleted repeat sequences compared to all repeat sequences from 293T, hESCs, and meso. (F) Bubble plot showing the expression of the rRNAs, LTR, LINE, and SINE classes in P-bodies, cyto, and whole cells of 293T, hESCs, and meso. Bubble sizes represent Z-scores normalized to sample, and colors indicate expression level. Z-score is normalized to sample by CPM (counts per million). (G) PCA plot showing the distribution of P-bodies, cyto, whole cells from 293T, hESCs, and meso for the expression of TEs. (H) Bubble plot showing the expression of the rRNA and TE families in P-bodies, cyto, and whole cells of 293T, hESCs, and meso. Bubble sizes represent Z-scores normalized to sample, and colors indicate expression levels normalized to sample by CPM. (I) Bar plot showing numbers of differentially expressed TEs in hESCs and meso, and P-body enriched or P-body depleted in 293T, hESC and meso. Colors indicate TE families. (J) Heatmap showing the expression of the representative TEs in P-bodies or cyto of 293T, hESCs, and meso. (K) Heatmap showing the expression of embryogenesis-specific TEs in P-bodies or cyto of 293T, hESCs, and meso. LTR_5Hs is specific for morula. LTR7Y is enriched in blastocyst. While LTR7 is highly expressed in blastocysts and primed hESCs. **See also Figure S6 and S7.**

The proportion of aligned mRNAs from P-bodies is generally lower than that from cytoplasm, and the number of protein-coding genes in P-bodies is lower compared with cytoplasm, especially in meso (Figure 5D, Figure S7A). Given that lncRNAs were also more enriched in cytoplasm, we then tested whether other non-coding RNAs, such as ribosomal RNAs (rRNAs) and transposable elements (TEs), were more enriched in P-bodies. Interestingly, rRNA are highly enriched in P-bodies, extremely in meso (Figures 5E-F, Figure S7A). The proportion of TEs in P-bodies is comparable to that in cyto of 293T cells and hESCs, but significantly lower in the P-bodies of meso (Figure 5E). PCA plot of TE families revealed that whole cells of hESCs are clustered with cyto but separated away from P-bodies, while whole cells of meso are more similar to the P-bodies (Figure 5G), suggesting the dynamics of P-body composition of TEs during hESC differentiation. This is in accordance with the reduction of differential expressed genes in meso. In particular, ERV1, LINE1 and ALU are significantly enriched in P-bodies of hESCs, while rRNA LSU and SSU are more enriched in P-bodies of meso (Figure 5H, Figure S7B). Differential expression analysis revealed that hESCs have a higher enrichment of TEs at the cellular level compared to meso (Figure 5I, Figures S7C-E), consistent with the finding during embryogenesis^36^. P-bodies of hESCs also shows a significant enrichment of TEs compared to 293T and meso (Figure 5I, Figures S7C-E, Table S4). The expression of TEs are dynamically regulated during early embryogenesis and different states of hESCs. LTR5_Hs is highly expressed in morula, LTR7Y is highly enriched in blastocysts, and LTR7 is highly expressed in blastocyst and primed hESC^37^. ERVs, LTR5_Hs, and LTR7Y are highly expressed in hESCs compared to meso, and particularly enriched in P-bodies of hESCs, which may imply their degradation during hESC to meso differentiation, similar to NANOG (Figure 5J- K). Taken together, non-coding RNAs are highly enriched in P-bodies, and P-bodies might be involved in the degradation of TEs during hESC to meso differentiation.

### P-bodies are involved in the regulation of hESC to mesoderm differentiation

Next, we aimed to detect the involvement of P-bodies in the regulation of hESCs. To test whether the loss of pluripotency during stem cell differentiation is associated with the changes of P-body composition, we treated hESCs with TGFβi or induced to meso for differentiation. We observed a decrease in the number of P-bodies with a concomitant downregulation of pluripotency markers in both TGFβi-triggered differentiation and meso differentiation (Figures 6A-B, Figure S8A). To further demonstrate the involvement of P-bodies in regulating hESC differentiation, we examined the effects of LSM14A over-expression in hESCs and in meso. Although LSM14A overexpression had marginal effect on the transcriptome of hESCs, it significantly affected the expression of about 300 genes in meso (Figures 6C-D, Table S5), with 40% of the upregulated differentially expressed genes being shared with hESCs or mesodermal cells, and 50% of the downregulated genes overlap with genes enriched in mesodermal cells (Figure 6E). Moreover, these DEGs are closely associate with developmental regulation (Figure S8B). These results suggest that P-bodies are involved in the regulation of hESC to meso differentiation.

**Figure 6.**
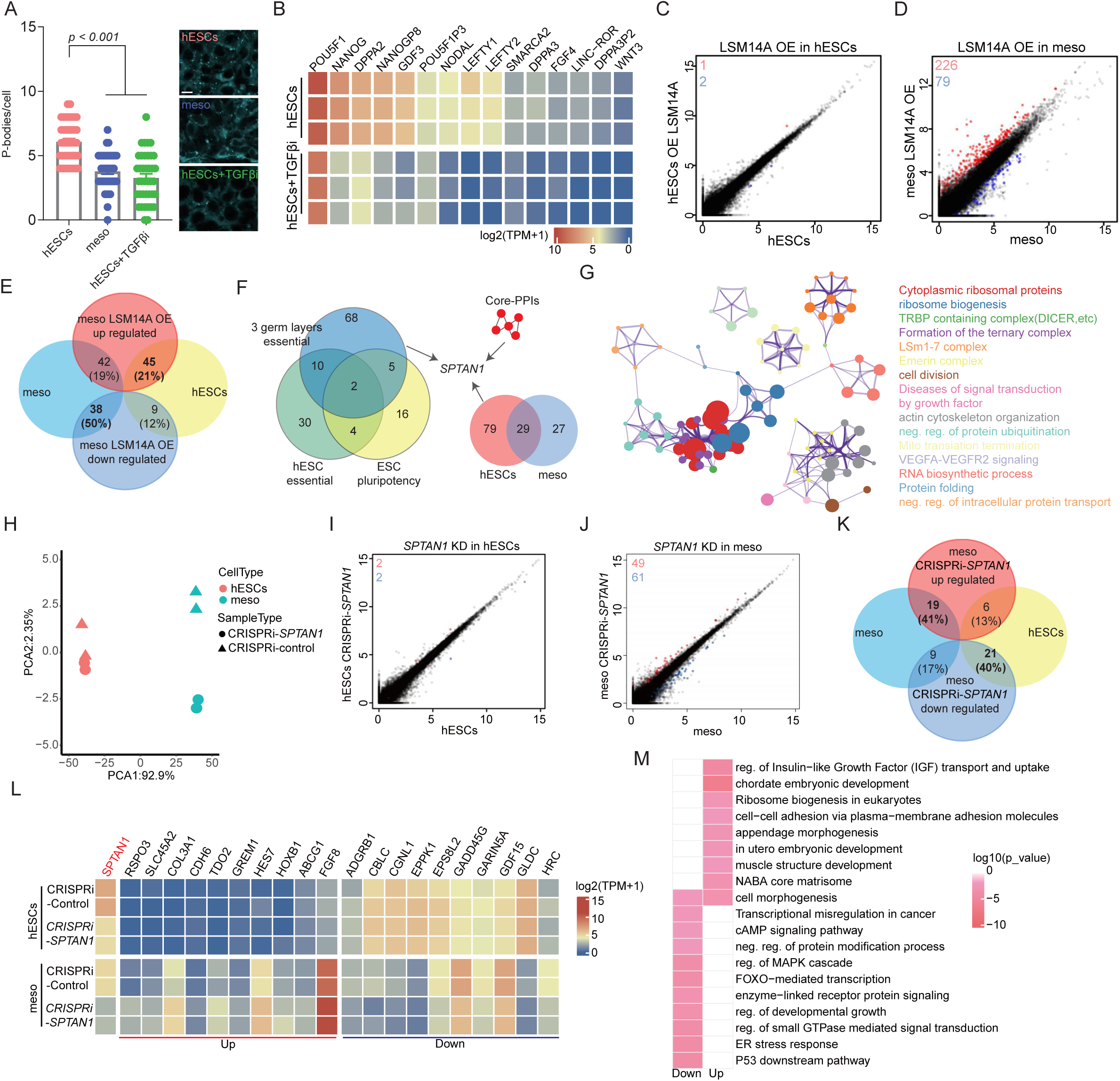
LSM14A and *SPTAN1* are involved in the regulation of hESC to mesoderm differentiation via P-bodies. (A) Quantification of P-bodies in hESCs, meso, and hESCs under TGFβ inhibitor treatment. DDX6-GFP-kncokin reporter line was used to visualize P-bodies. N = 40 cells for each group. Data is shown as mean ± SEM. The numbers of P-bodies per cell are: 6.08 ± 0.25 for hESCs; 53.8 ± 0.23 for meso; 3.28 ± 0.3 for hESCs+TGFβi (hESCs treated with TGFβi). Scale bar:10μm. (B) RNA-seq analysis showing the expression of the selected pluripotency genes in hESCs and hESCs+TGFβi. (C-D) Scatter plots showing the comparisons of the DEGs in hESCs with over-expression of LSM14A to control hESCs (C), and DEGs in meso with over-expression of LSM14A to control meso (D). (E) Venn plot showing the overlap of up-regulated and down-regulated genes upon over-expression of LSM14A to hESC-and meso-enriched genes. (F) Venn plot showing the overlap of P-body enriched genes from hESCs and meso with hESC essential genes^38^, hESC differentiation essential genes^39^, and pluripotency essential genes^40–42,53^. There are 135 genes from the overlapped fraction in total, and *SPTAN1* is one of the representative genes. (G) Functional clusters of the 135 genes from (F). (H) PCA plot showing the RNA-seq samples of control and CRISPRi-*SPTAN1* hESCs and meso. (I-J) Scatter plots showing the comparisons of the DEGs in hESCs (I) and meso (J) for control and CRISPRi-*SPTAN1*. (K) Venn plot showing the overlap of up-regulated and down-regulated genes upon CRISPRi-*SPTAN1* to hESC-and meso-enriched genes. (L) Heatmap showing the hESC-and meso-enriched genes from CRISPRi-*SPTAN1* meso to control meso. *SPTAN1* is highlighted in red. (M) Heatmap showing the functional enrichment of DEGs from CRISPRi-*SPTAN1* meso to control meso. **See also Figure S8.**

To identify the potential P-body enriched genes that are involved in regulating hESC differentiation, we compared P-body enriched mRNAs of hESCs and meso to the published data including: (1) essential genes for hESC maintenance^38^, (2) essential genes for hESC^39^, and (3) essential genes for ESC pluripotency^40–42^. A total of 135 genes were identified (Figure 6E, table S6), including *SPTAN1* (*spectrin alpha 1*), a member of the spectrin family of filamentous cytoskeletal proteins that serve as essential scaffold proteins. *SPTAN1* encodes an alpha spectrin specifically expressed in nonerythrocytic cells, and involved in various cellular functions, including DNA repair and cell cycle regulation. Mutations in SPTAN1 have been discovered in diseases such as early infantile epileptic encephalopathy^43^. *SPTAN1* is also the core gene of cytoskeleton terms and PPIs (Figures 3E-H, Figure 6F). These observations promoted us to test the involvement of *SPTAN1* in hESC to meso differentiation. To achieve this, we constructed *SPTAN1* knockdown (KD) cell lines by CRISPRi, verified by gene track of RNA-seq and qPCR (Figures S8D-E). Importantly *SPTAN1* KD did not affect the expression of pluripotency genes in hESCs (Figures S8F-H), of the meso markers in meso (Figures S8G, I). We then investigate the influence on RNA transcripts. *SPTAN1* KD cells did not differ from the control group in hESCs, but were divided into two populations in meso by PCA analysis. Importantly, *SPTAN1* KD had only 4 DEGs (including *SPTAN1*) in hESCs (Figure 6I), but significantly perturbed 110 genes expression in meso (Figure 6J, Table S7), with 19 genes (41%) of the upregulated differentially expressed genes being shared with meso, and 21 genes (40%) of the downregulated genes overlap with genes enriched in hESCs (Figure 6K, L). Moreover, the upregulated genes are enriched in the terms including cell-cell adhesion, embryonic development, morphogenesis, which is very similar with the functions of P-bodies deleted genes (Figure 6M, Figures S5C-E). While the downregulated genes are enriched in the terms including cAMP signaling, GTPase and protein modification, which is found in the terms of P-body enriched genes of hESCs or meso (Figure 6M, Figure 3G, Figure S4D). Taken together, these results suggest that the P-body enriched gene *SPTAN1* is involved in hESC to meso differentiation.

## DISCUSSION

In this study, we characterized the RNA profiles of the P-bodies from 293T cells, hESCs, and hESC-derived mesodermal cells. We discovered the common features of RNAs in P-bodies compared with RNAs from other cytoplasmic compartments such as cytosol, mitochondria, and stress granule. The RNAs in P-bodies contain lower GC contents, which may be related to the reduced RNA stability and facilitate the formation of unstructured regions, contributing to the formation of P-bodies. In contrast, conserved protein-coding genes tend to have higher GC content, consistent with the requirement for structural stability. We also found lower m^6^A modification on the mRNAs in P-bodies. This may imply the reduced RNA processing complexity with respect to translation and degradation processes, as m^6^A modification could modulate the secondary structures of RNAs, which in turn will establish the binding or recognition sites for RNA-protein interactions^44^.

Our results reveal the differences of P-body components among different cell types (Figure 3). Cytoplasmic mRNAs are mainly associated with basic cellular processes and conserved functions, while in contrast, P-body enriched mRNAs and lncRNAs show cell-specific features. Importantly, P-body enriched genes and regulatory factors are significantly reduced during hESC to meso differentiation, and the P-body enriched genes in meso are mainly associated with translation regulation. A previous study highlighted the influence of the endoplasmic reticulum (ER) and translation on P-body abundance, revealing the negative correlation between cellular translation levels and P-body abundance^45^. During hESC to meso differentiation, we observed the loss of regulatory components and the accumulation of translation-related genes in P-bodies, which may correspond to the reduced number of P-bodies. Furthermore, translational regulation is a key feature of embryonic development^46^, further emphasizing the potential critical roles that P-bodies play during embryogenesis. We also derived common diseases associated genes from OMIM, and found that most of these genes are excluded from P-bodies. This is consistent with our findings, because P-body depleted genes are mainly associated with basic cellular functions and development. However, the disease associated P-bodies enriched genes should be noted, as most of these genes are less expressed at cellular level, indicating that these genes are regulated or degraded but also important, improper express of which are associated with diseases.

Our analysis of lncRNAs has revealed their dual roles (Figure 5). P-body-enriched lncRNAs are primarily involved in functions related to RNA metabolism, degradation and regulatory processes such as alternative splicing and modifications. Cytoplasmic lncRNAs, on the other hand, can directly interact with proteins and are involved in post-transcriptional regulation, RNP complex formation and protein scaffolding^34,47^. Notably, hESCs exhibit more protein-lncRNA interactions compared with 293T cells or meso, indicating the distinct role of P-bodies in hESCs. LncRNA expression is partially regulated by adjacent transposable elements (TEs) when they become transcriptionally activated during embryogenesis, but are silenced gradually during development^48,49^. Furthermore, the expression of different families of LTRs are highly correlated with different stages of early embryogenesis. Interestingly, P-bodies in hESCs significantly enrich different types of TEs compared with meso (Figure 5). Our study also suggests the potential roles of P-bodies in regulating the differential

We also observed the relationship between P-body quantity and the differentiation of stem cells. Differentiation of hESCs leads to a relatively reduced number of P-bodies. Interestingly, overexpression of LSM14A or *SPTAN1* knockdown does not impair the transcriptome of hESCs, but disrupts the expression of development-associated genes during mesoderm differentiation (Figure 6). Importantly, LSM14A OE upregulated genes and *SPTAN1* KD downregulated genes have more overlap with hESC-specific genes, and conversely, LSM14A OE downregulated genes and *SPTAN1* KD upregulated genes have more overlap with meso-specific genes, most of these genes are involved in cell development (Figures 6). These results indicate that P-bodies may function as the storage sites for RNAs in hESCs. These RNAs will be released from P-bodies to execute their functions during hESC differentiation. Altogether, our results suggest the involvement of P-bodies in the regulation of stem cell differentiation.

## ACKNOWLEDGEMENT

The pC13N-dCas9-BFP-KRAB and CLYBL-L/R plasmids were kindly provided by Prof. Ruilin Tian (Southern University of Science and Technology, China), the pLenti-U6-gRNA-EF1-mCherry plasmid was kindly provided by Prof. Chaochen Wang (Zhejiang University, China). We thank Ms. Hangchen Shen from the core facility of of Zhejiang University-University of Edinburgh Institute (Zhejiang University, China) for technical assistance in establishing the P-body purification system in this study.

This study was supported by “Chaoyong” Special Support for Postdoctoral Researchers at International Campus, Zhejiang University (to J.J.), the National Natural Science Foundation of China (32270835 to D.C.), and Zhejiang Natural Science Foundation Key Program (Z22C129553 to D.C.).

## AUTHOR CONTRIBUTIONS

J.J. and D.C. conceived, initiated, and designed the project; J.J. performed most of the experiments and analyzed most of the data Analysis. Q.S., X.X., R.G. generated cell lines used in this study and performed most molecular cloning and genotyping works; S.X. helped with bulk RNA-seq data analysis, M.J. helped with data interpretation and preparation of the manuscript; J.J. and D.C. wrote the manuscript. All authors edited and approved the manuscript.

## DECLARATION OF INTERESTS

The authors declare no competing interests.

## SUPPLEMENTARY FIGURES AND LEGENDS

**Figure S1.**
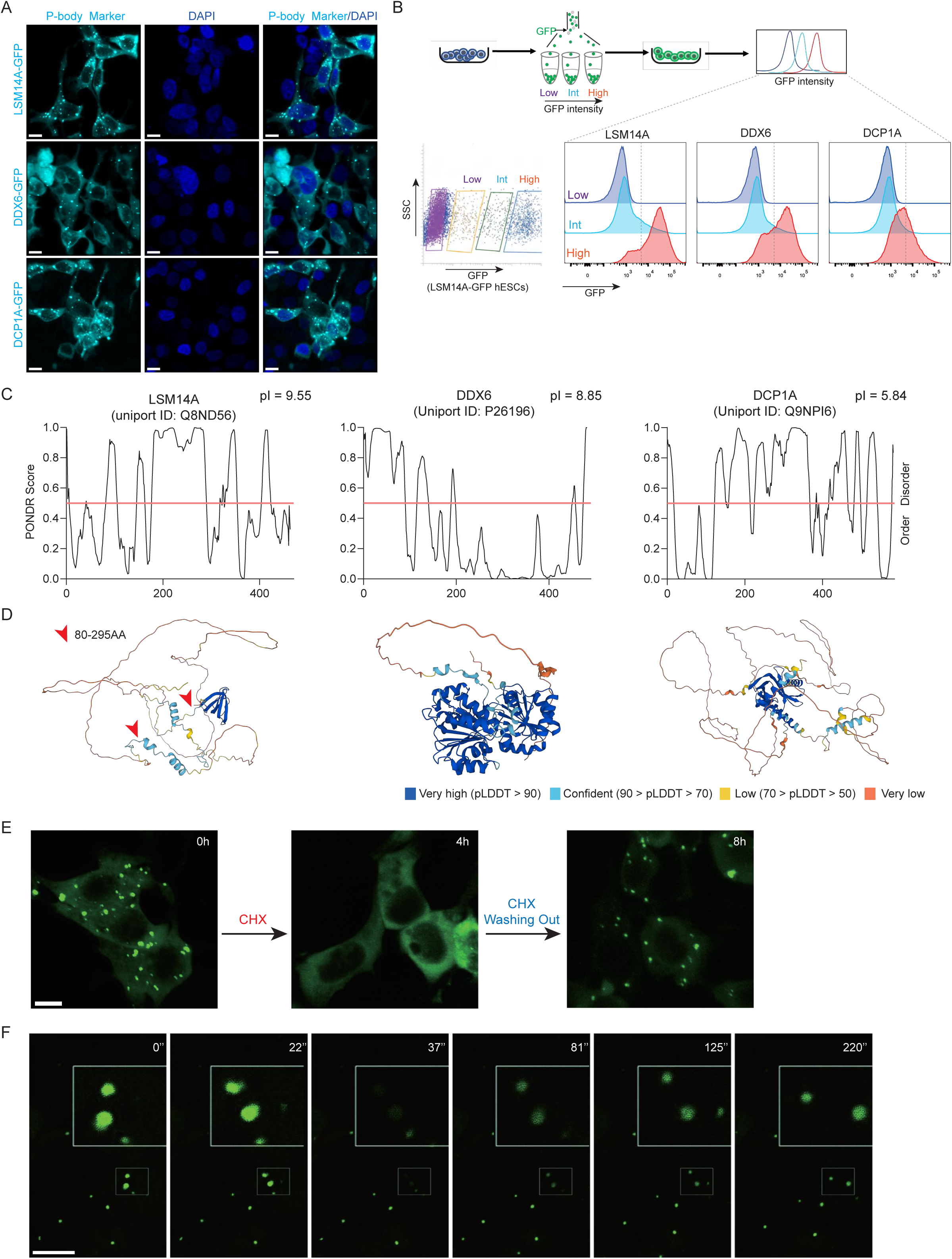
Screening of reporter lines for P-body purification. Related to Figure 1. (A) Labelling of P-bodies with canonical P-body markers LSM14A, DDX6 and DCP1A fused with GFP in 293T cells. Scale bar: 10μm. (B) Persistency test of the P-body reporter lines. GFP positive cells were sorted to three groups based on the high, intermediate (int), and low fluorescent intensity followed by 3 days of culture to test for GFP intensity. (C-D) Disordered characteristics, theoretical isoelectric points, and predicted structures of DCP1A, DDX6, and LSM14A proteins analyzed by PONDR, Expasy, and Alphafold2, respectively. pI: theoretical isoelectric point. (E) Cycloheximide (CHX) treatment of 293T cells for P-body dynamics. Scale bar: 10μm. (F) P-body dynamics test by fluorescence recovery after photo bleaching (FRAP) in 293T cells. Scale bar: 10μm.

**Figure S2.**
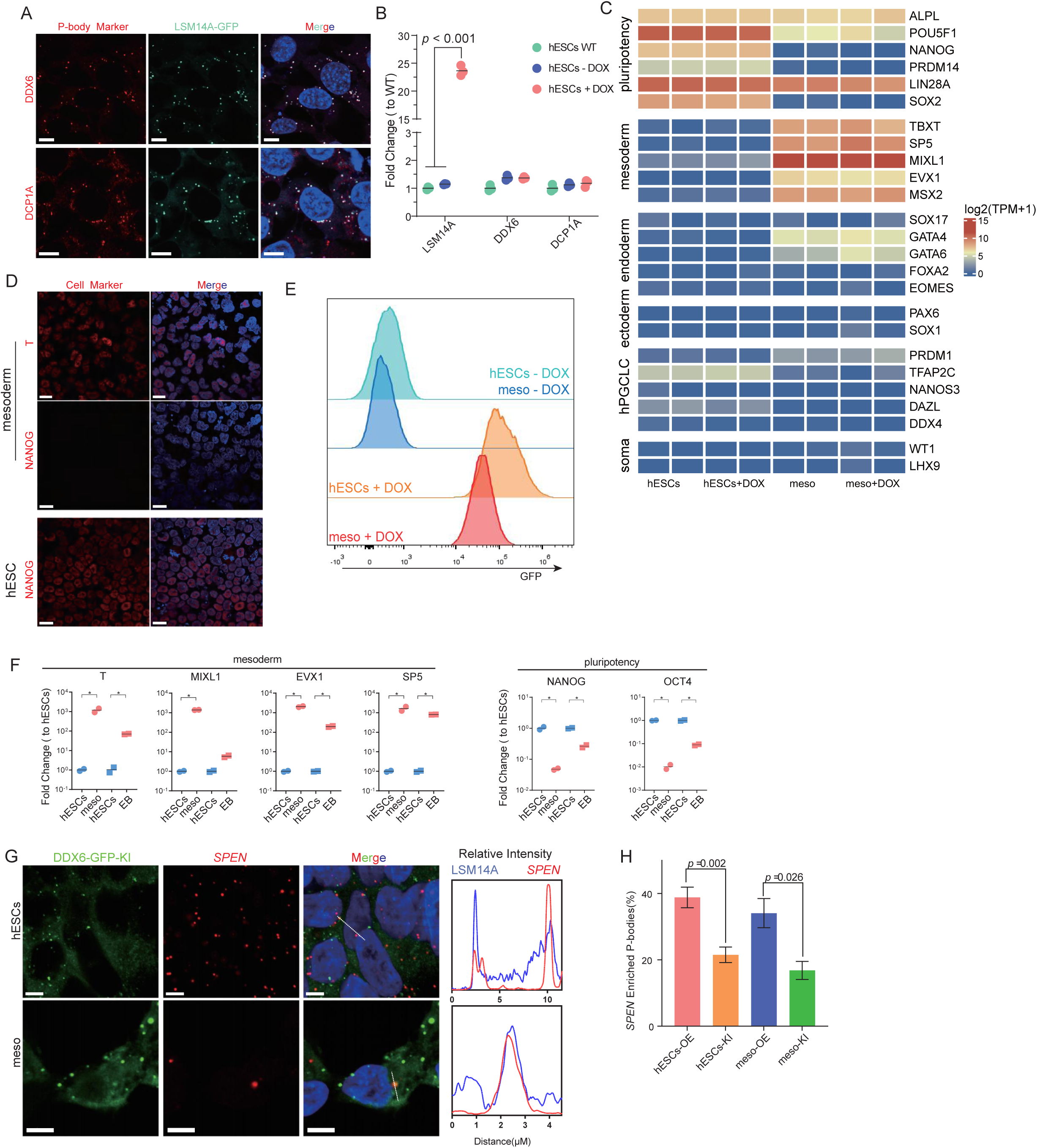
Purification of P-bodies using tetON-LSM14A-GFP reporter line. Related to Figure 1. (A) Immunofluorescence of DDX6 and DCP1A in hESCs with LSM14A-GFP reporter 4 days after doxycycline (Dox) induction. Scale bar: 10 μm. (B) qPCR verification of the effect of ectopic expression of *LSM14A* on the expression other P-body markers *DDX6* and *DCP1A*. (C) RNA-seq analysis showing the expression of pluripotency and differentiation-related genes in hESCs and meso with or without Dox for over-expressing *LSM14A-GFP*. (D) Immunofluorescence of T(BRACHYURY) and NANOG in hESCs and meso after Dox induction of *LAM14A-GFP*. DAPI (blue) is counterstained to indicate nuclei. Scale bar: 15μm. (E) Flow cytometry showing the distribution of GFP positive cells in hESCs and meso with or without Dox induction. (F) qPCR detection of the expression of mesoderm and pluripotency markers in hESCs, embryonic bodies (EB), and meso. (G) RNAscope images showing the enrichment of *SPEN* mRNAs (red) in P-bodies from *DDX6-GFP* knockin (KI) hESCs and meso. DAPI is counterstained to indicate nuclei. Scale bar: 5 μm. (H) The percentages of P-bodies containing *SPEN* mRNAs in hESCs with *LSM14-GFP* over-expression (OE) and hESCs with *DDX6-GFP* knockin (KI) (n = 40 cells, data is shown for mean±SEM. hESC-OE: 38.8±3.1; hESC-KI: 21.5±2.3; meso-OE: 37±4.4; meso-KI:16.8±2.7).

**Figure S3.**
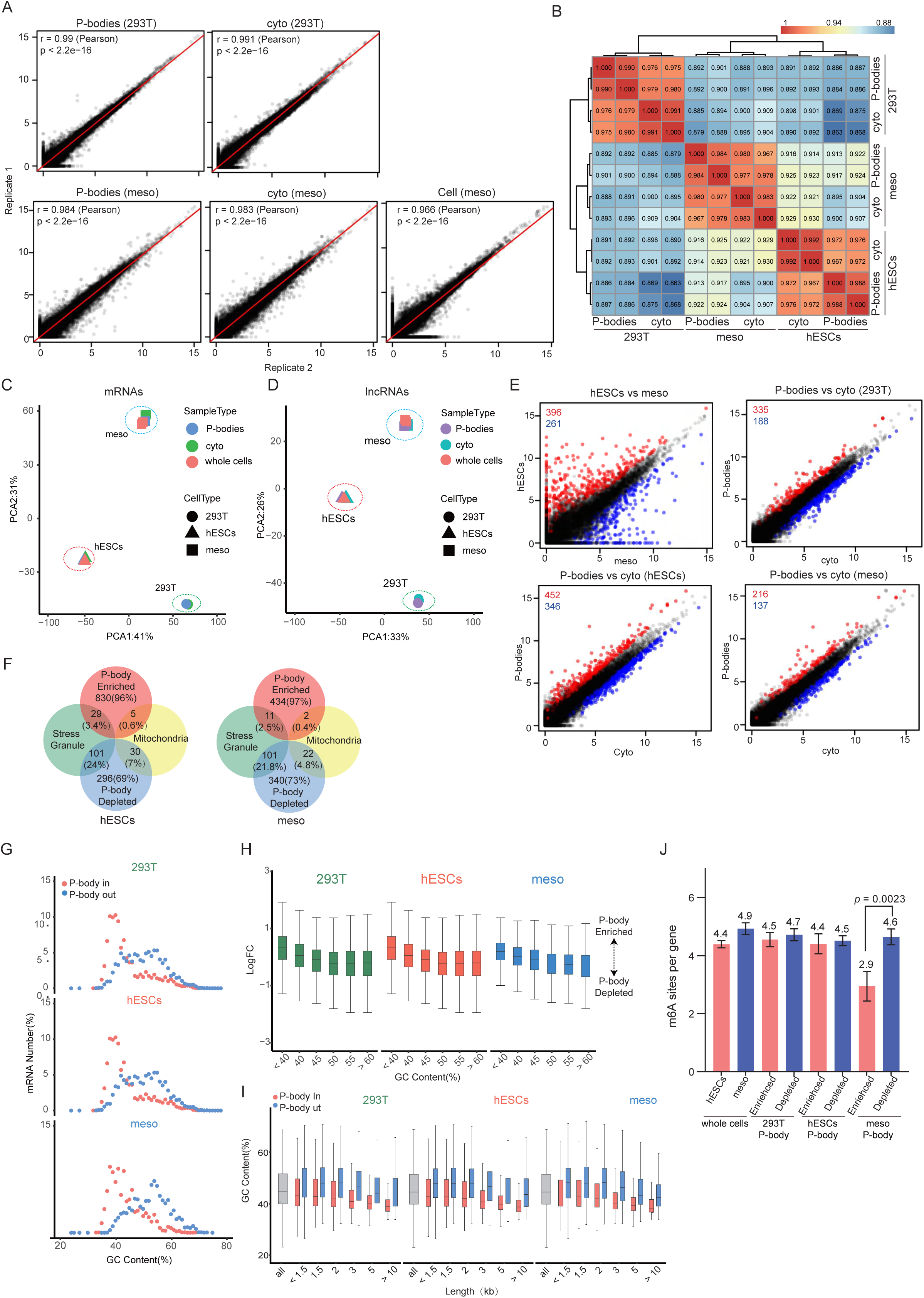
Analysis of the features of P-body enriched and P-body depleted genes. Related to Figure 2. (A) Scatter plots showing the correlation of the two biological replicates of transcriptomes of P-bodies from 293T and meso, cyto from 293T and meso, and whole cells from meso. (B) Correlation analysis for the transcriptomes of P-body and cyto from 293T, hESCs, and meso. (C-D) PCA plot showing the mRNA (C) or lncRNAs (D) of the whole cells, cyto, and P-bodies from 293T, hESCs, and meso. (E) Scatter plots showing the DEGs for hESCs vs meso, P-body vs cyto in 293T, hESCs, and meso. (F) Venn plots showing the comparison of P-body enriched genes and P-body depleted genes from hESCs and meso, with genes enriched for stress granules^50^ and mitochondria^51,52^. (G) The distribution of GC contents for P-body in and P-body out genes in 293T, hESCs, and meso. The median of GC contents of P-body in vs P-body out for each cell types are 40.38% vs 49.5% for 293T, 40.8% vs 46.0% for hESCs, and 41.5% vs 52.36% for meso. (H) The distribution of GC contents for P-body enriched and P-body depleted mRNAs in 293T, hESCs, and meso. (I) The GC contents of P-body in (FC>0) and P-body out (FC<0) genes for transcripts with different lengths. (J) Bar plots showing the numbers of m^6^A site per P-body enriched gene or per P-body depleted gene in 293T, hESCs, and meso.

**Figure S4.**
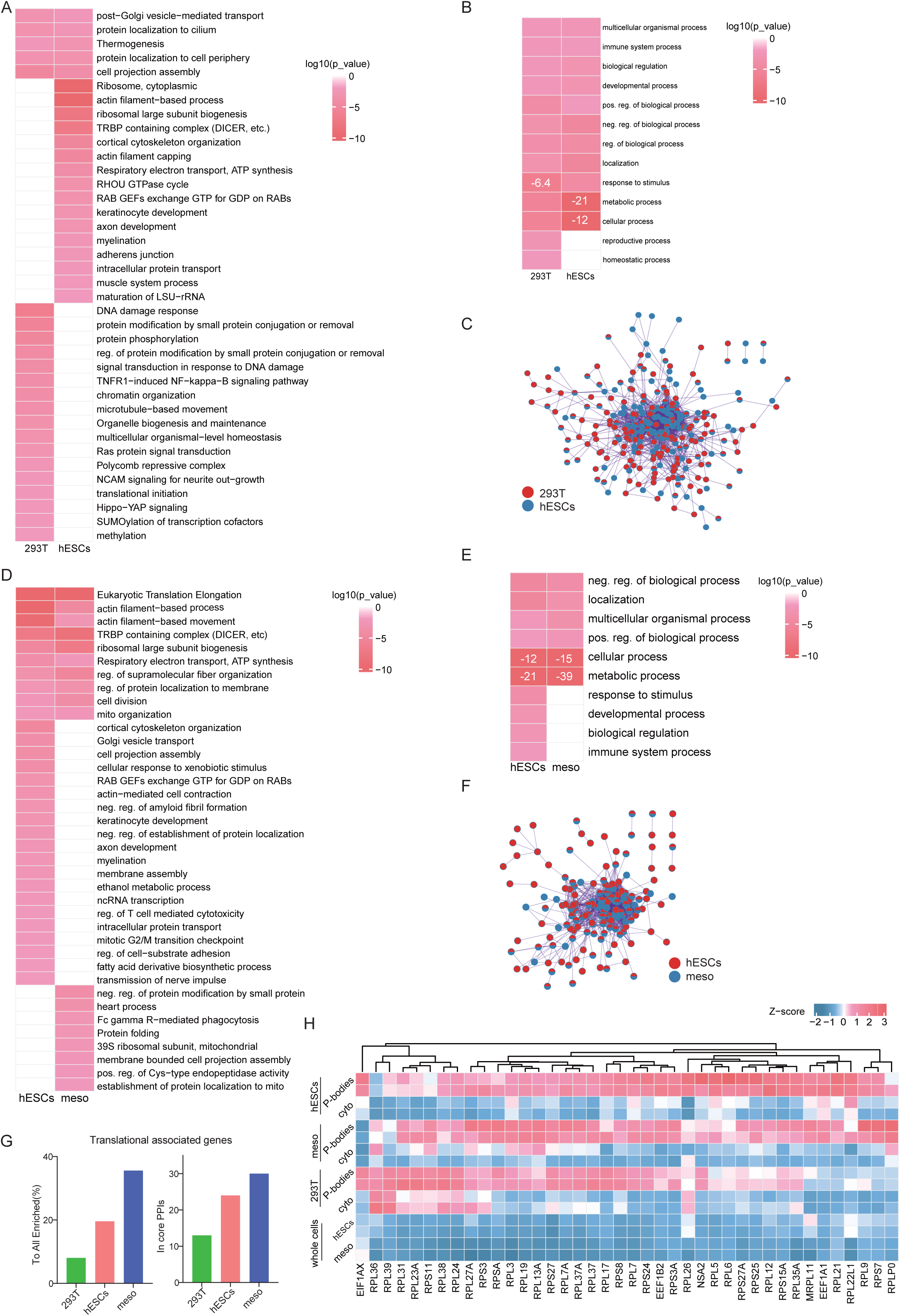
Functional clustering and protein-protein interactions of P-body enriched genes. Related to figure 3. (A) Heatmap showing functional enrichment of P-body enriched genes of 293T and hESCs. Related to Figure 3E. (B) Heatmap showing the parental Gene Ontology enrichment of P-body enriched genes of 293T and hESCs. Related to Figure 3E. (C) PPI network of all protein coding genes showing the interactions in 293T (red) and hESCs (blue). Related to Figure 3F. (D) Heatmap showing functional enrichment of P-body enriched genes of hESCs and meso. Related to Figure 3G. (E) Heatmap showing the parental Gene Ontology enrichment of P-body enriched genes of hESCs and meso. Related to Figure 3G. (F) PPI network of all protein coding genes showing the interactions of P-body enriched in hESCs (red) and meso (blue). Related to Figure 3H. (G) Bar plots showing the percentage of translation-associated genes enriched in P-body to all genes (left) or to genes of PPI network (right) in 293T, hESCs, and meso (left). Related to Figures 3F, H. (H) Heatmap showing the translational associate genes of the core PPI network. Related to Figure S4G.

**Figure S5.**
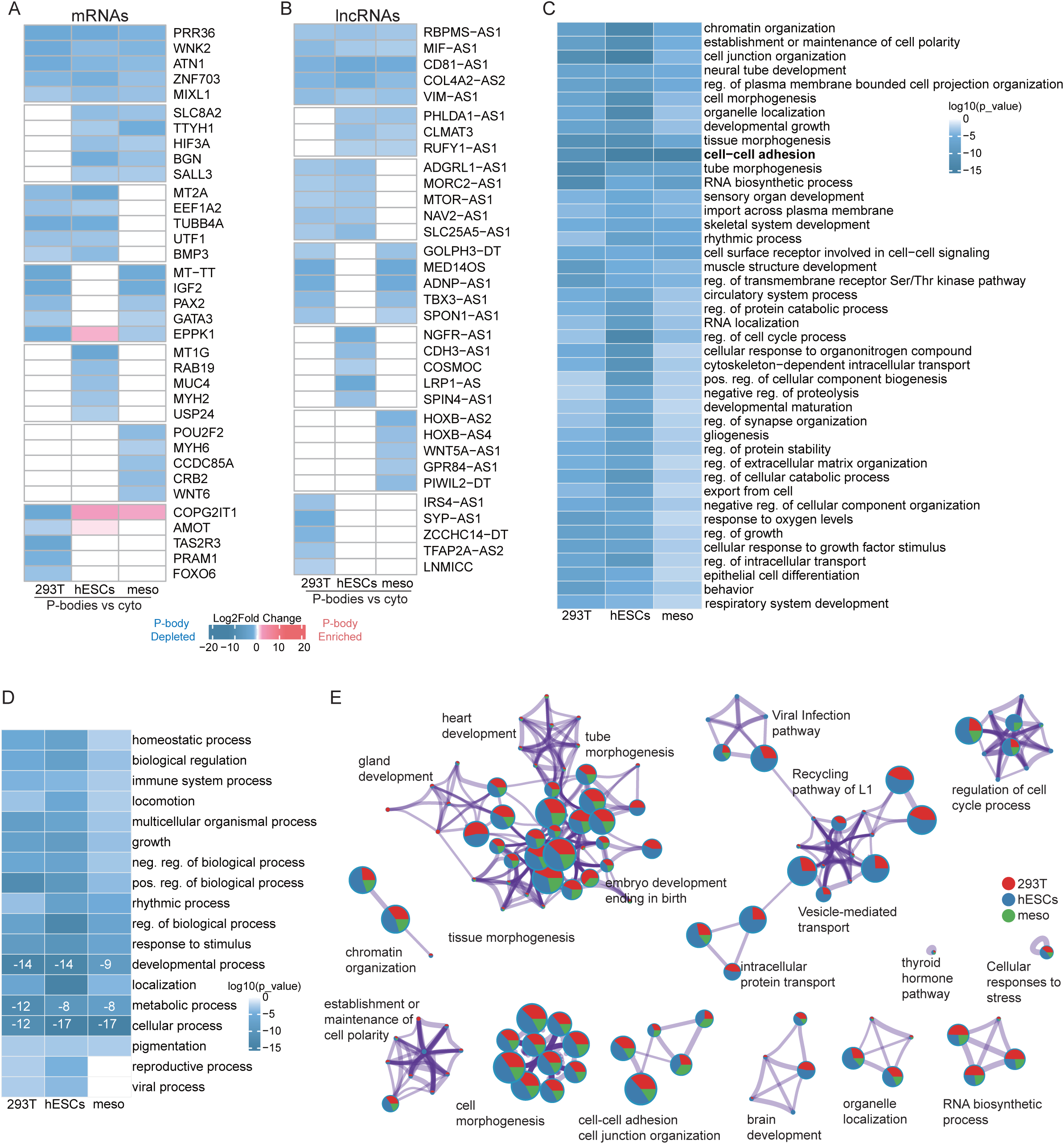
Characterization of the P-body depleted genes. Related to figure 3. (A) Heatmap showing the P-body depleted mRNAs shared or unique for different cell type(s) of 293T, hESC and meso. (B) Heatmap showing the P-body depleted lncRNAs shared or unique for different cell type(s) of 293T, hESC and meso. (C) Heatmap showing functional enrichment of P-body depleted genes from 293T, hESCs and meso. (D) Heatmap showing the parental Gene Ontology enrichment of P-body depleted genes from 293T, hESCs, and meso. (E) Functional Clusters showing the properties of P-body depleted genes from 293T (red), hESCs (blue), and meso (green).

**Figure S6.**
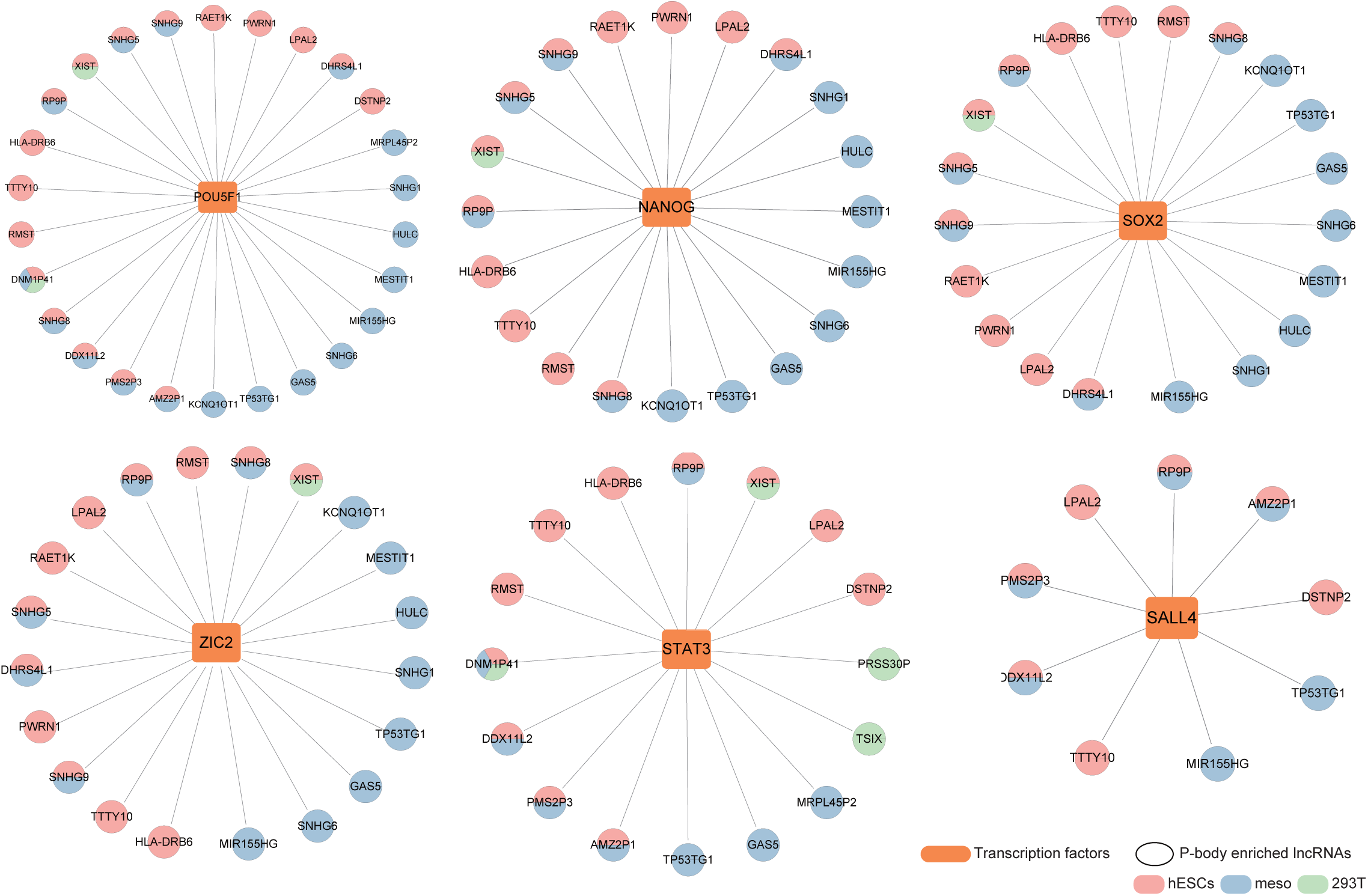
The interaction of Transcription factors with lncRNAs enriched in P-bodies. Related to figure 4. Interaction networks showing the pluripotency-related transcription factors and associated lncRNAs enriched in P-bodies from 293T, hESCs, or meso based on lncSEA (https://bio.liclab.net/LncSEA/index.php).

**Figure S7.**
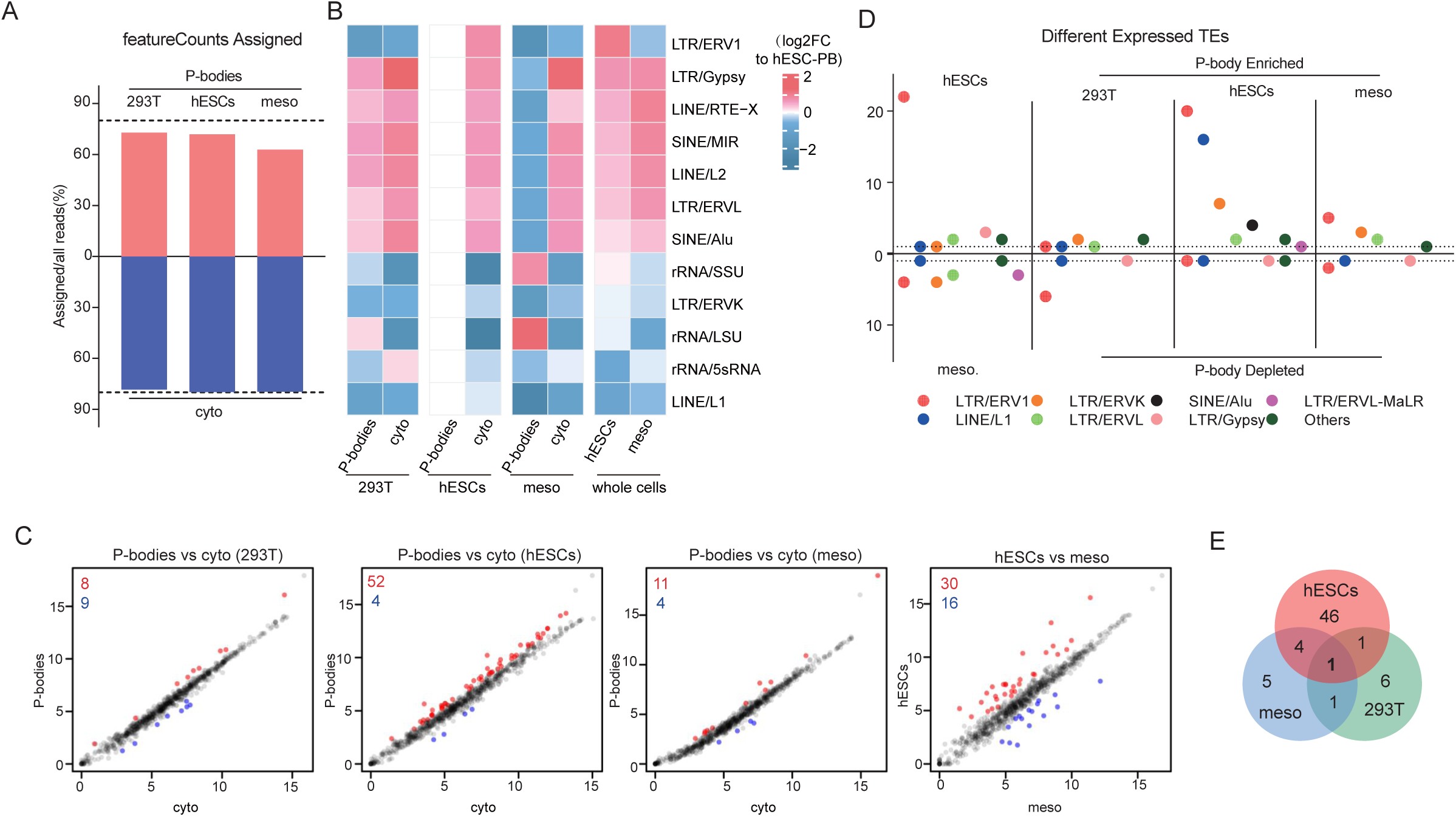
The expression of rRNA and TEs in P-bodies. Related to figure 5. (A) Bar plot showing the percentages of assigned transcripts to all transcripts of mRNAs from P-bodies and cyto of 293T, hESCs and meso by featureCounts. (B) Heatmap showing the expression of TEs and rRNAs in P-bodies and cyto from 293T, hESCs, and meso. (C) Scatter plot showing the enriched (red) and depleted (blue) TEs in P-bodies from 293T, hESCs and meso. (D) Dot plot showing the P-body enriched and P-body depleted TEs in 293T, hESCs and meso. Related to figure 6F. (E) Venn plot showing the overlaps of P-body enriched TEs from the 293T, hESCs, and meso.

**Figure S8.**
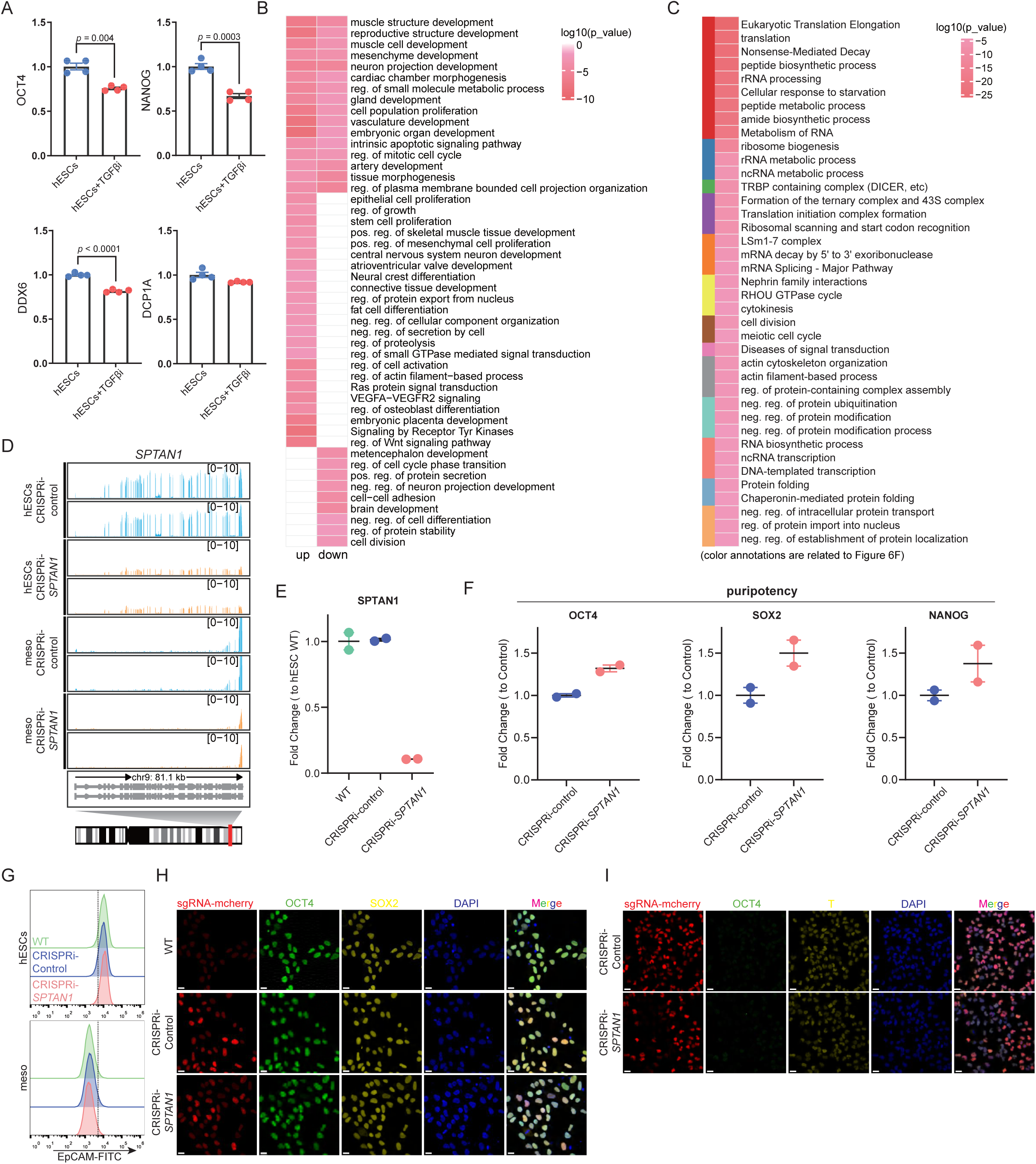
P-bodies might be involved in the hESC to mesoderm differentiation. Related to figure 6. (A) qPCR determination of the expression of the pluripotency-related genes (*OCT4*, *NANOG*) and P-body-related genes (*DDX6*, *DCP1A*) in hESCs and hESCs+TGFβi. (B) Heatmap showing the functional enrichment of DEGs from meso with over-expression of LSM14A to control meso. (C) Heatmap showing the functional enrichment of the 135 genes from Figure 6E. (D) Tracks showing the expression of *SPTAN1* in control and CRISPRi-*SPTAN1* hESCs and meso. (E) qPCR determination of the expression of *SPTAN1* in CRISPRi-*SPTAN1* and control hESCs. (F) qPCR determination of the expression of pluripotency-related genes (*OCT4*, *NANOG*) in control and CRISPRi-*SPTAN1* hESCs. (G) Flow cytometry showing the expression of EpCAM in hESCs and meso of wildtype, CRISPRi-control, and CRISPRi-*SPTAN1* cells. (H) Immunofluorescence showing the expression of mCherry (sgRNA, red), OCT4 (green), SOX2 (yellow) in hESCs of wildtype, CRISPRi-control and CRISPRi- *SPTAN1* cell lines. DAPI (blue) is counterstained to indicate nuclei. Scale bar: 20μm. (I) Immunofluorescence showing the expression of mCherry (sgRNA, red), OCT4 (green), T(yellow) in meso of CRISPRi-control and CRISPRi-*SPTAN1* cells. DAPI (blue) is counterstained to indicate nuclei. Scale bar:20μm.

## SUPPLEMENTARY TABLES

**Table S1.** Differentially expressed genes between P-bodies and cytoplasm across different cell types, related to Figure 2

**Table S2.** Differentially expressed lncRNAs between P-bodies and cytoplasm across different cell types, related to Figure 2

**Table S3.** Diseases related genes from OMIM expression between P-bodies and cytoplasm across different cell types, related to Figure 4

**Table S4.** Differentially expressed transposable elements and rRNAs between P-bodies and cytoplasm of 293T between P-bodies and cytoplasm across different cell types, related to Figure 5

**Table S5.** Differentially expressed genes between LSM14 overexpression and control in hESCs and mesodermal cells, related to Figure 6

**Table S6.** The screened 135 P-bodies enriched genes from hESCs or mesodermal cells related to ESC pluripotency, maintenance or differentiation, related to Figure 6

**Table S7.** Differentially expressed genes between SPTAN1 Knockdown and control in hESCs and mesodermal cells, related to Figure 6

## METHODS

### Key Resources Table

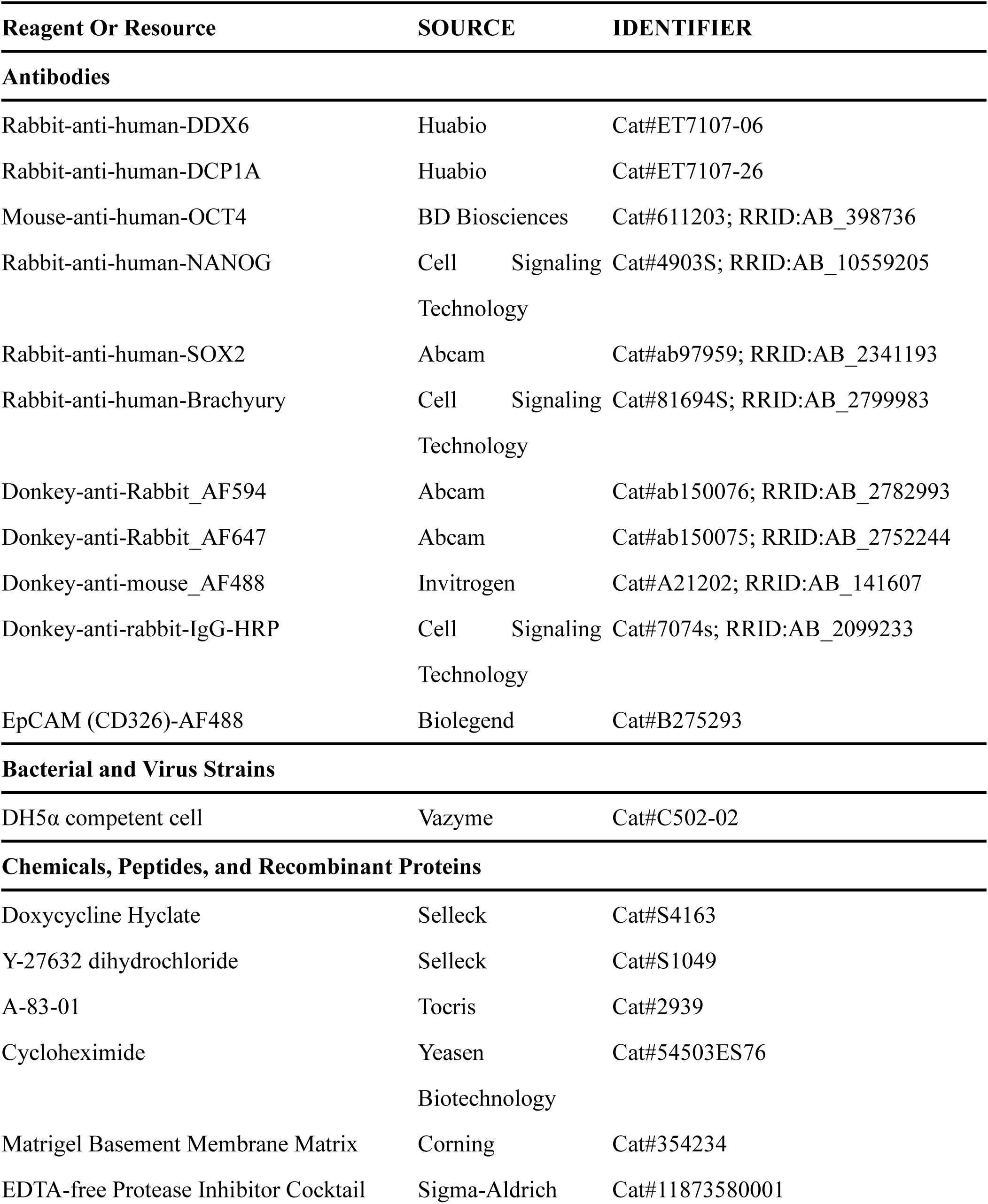
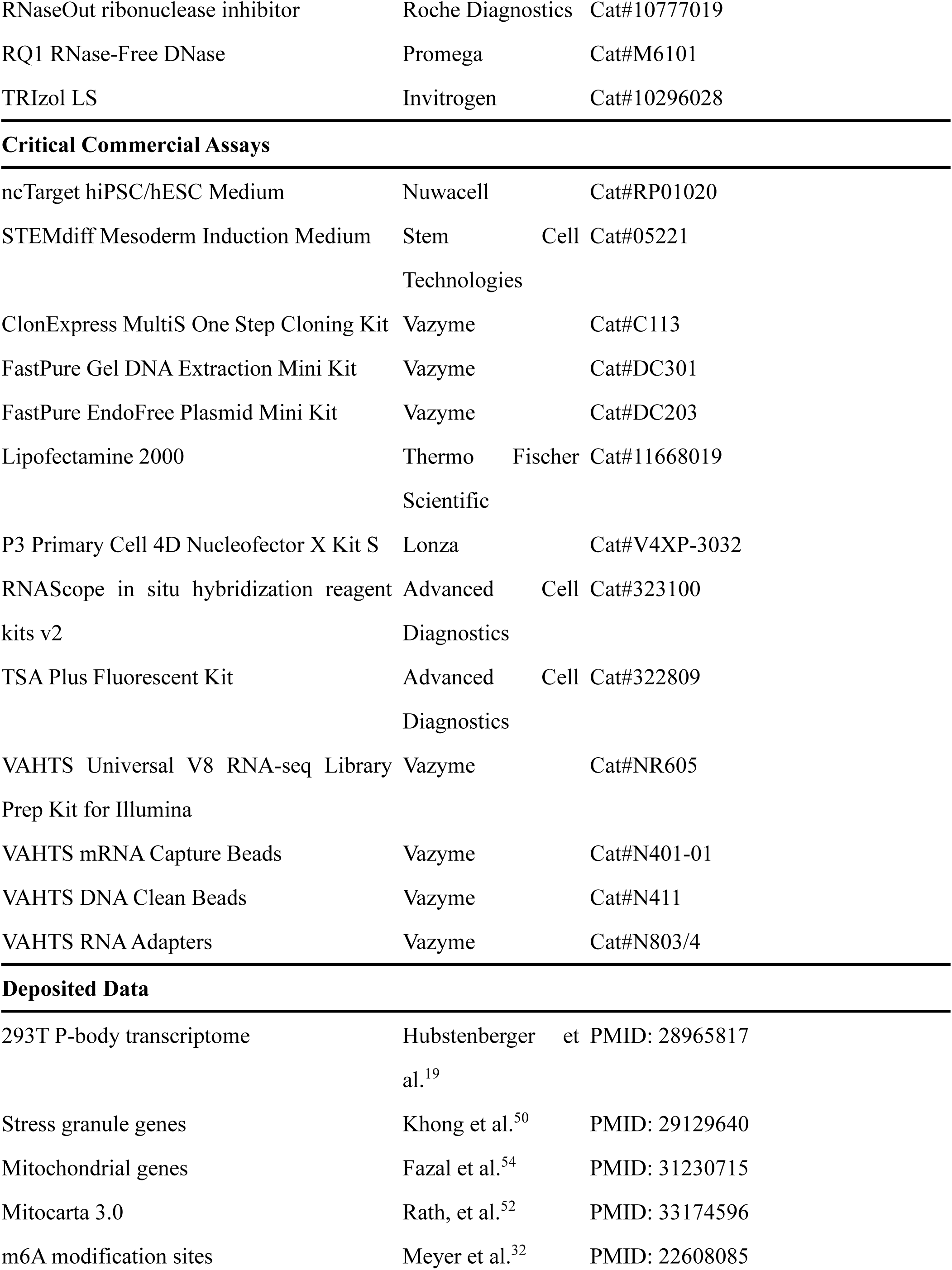
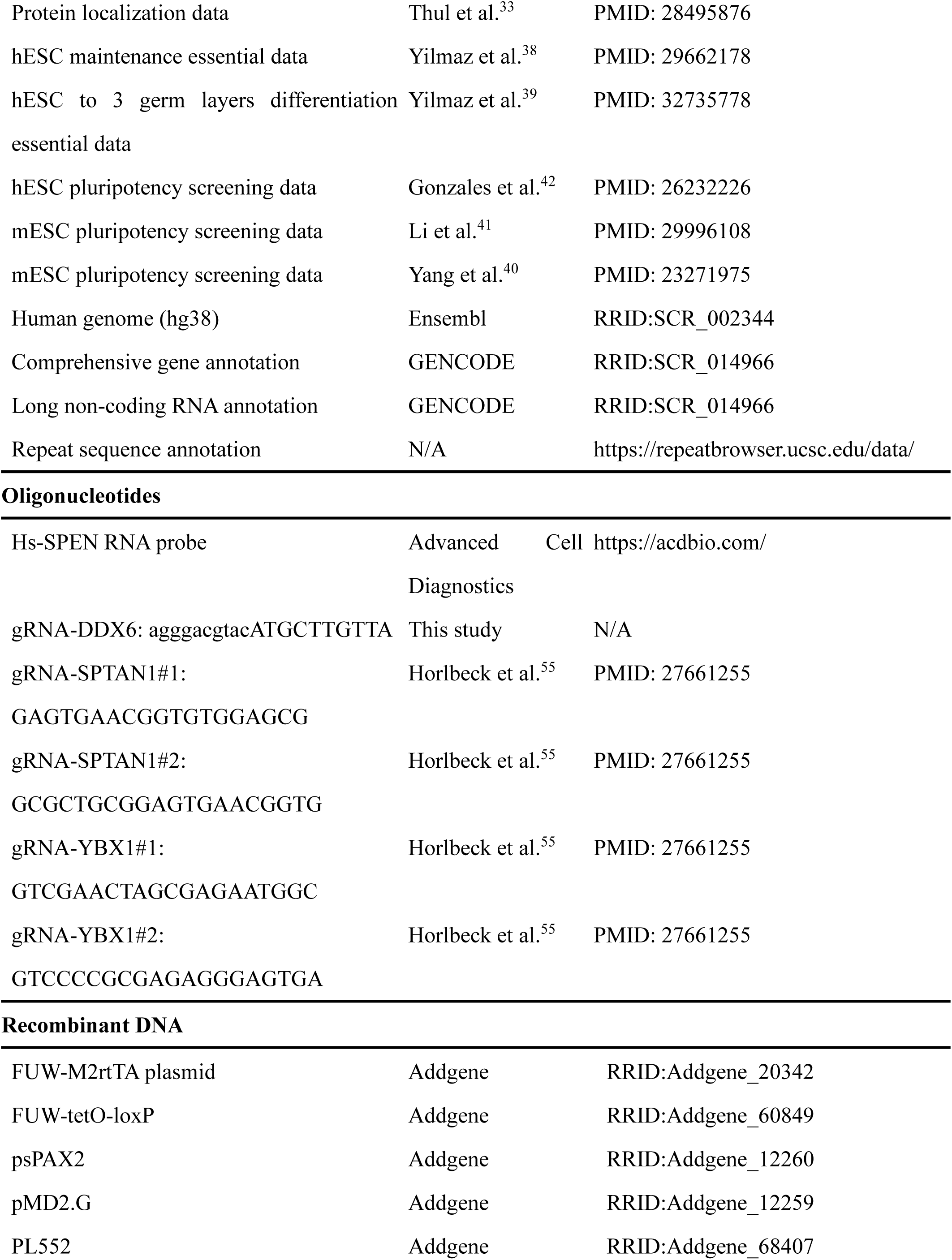
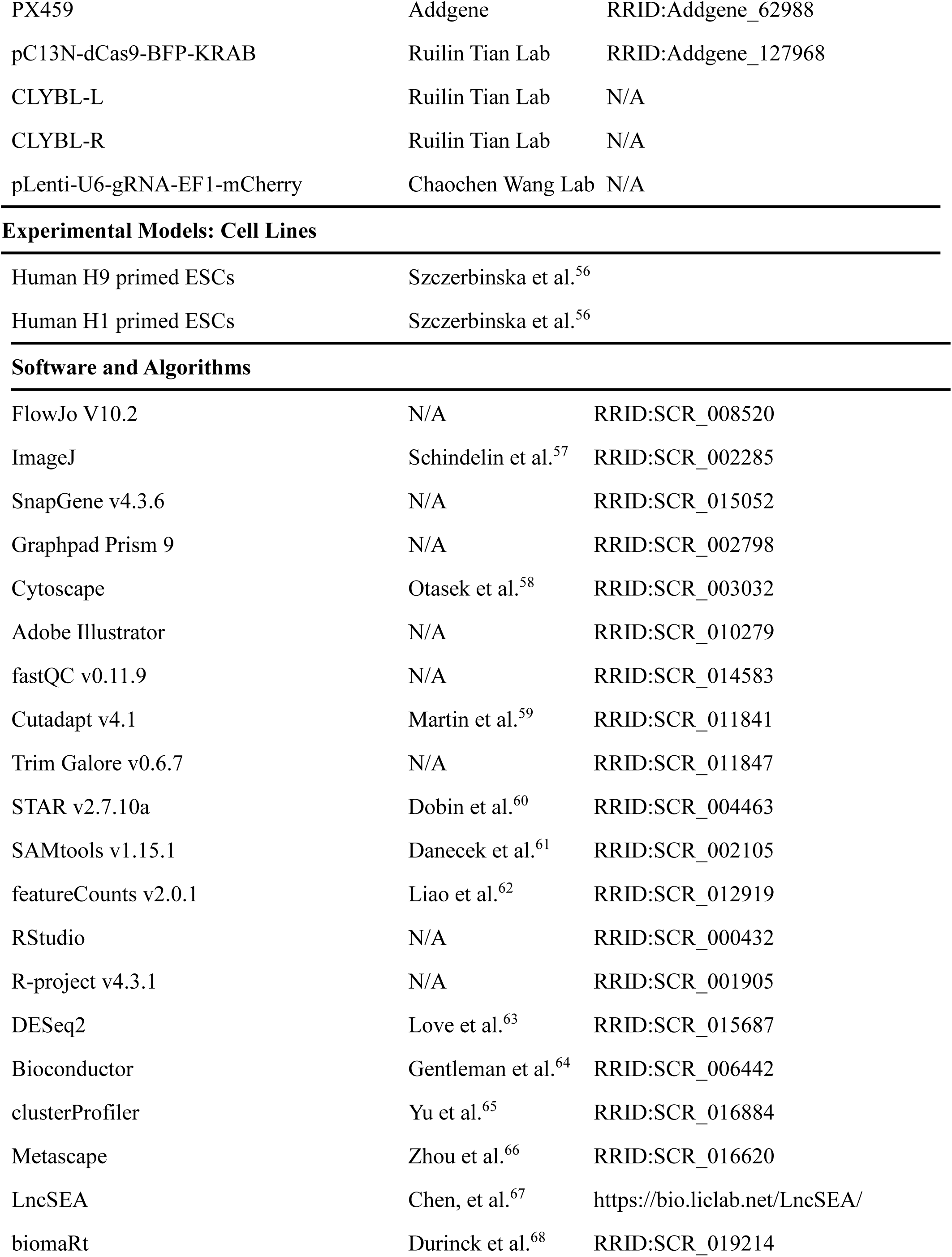
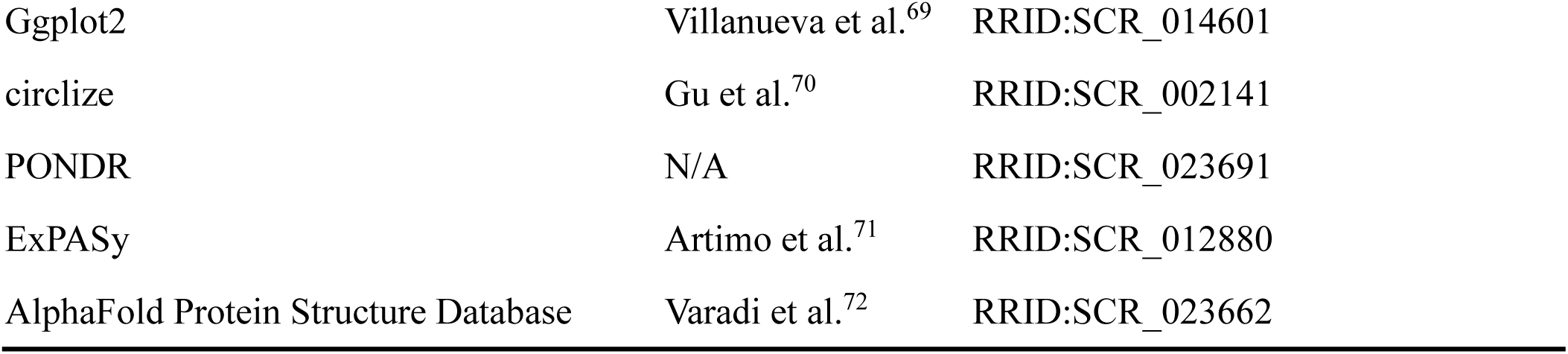

### Lead Contact and Materials Availability

Further information and requests for resources and reagents should be directed to and will be fulfilled by the Lead Contact, Di Chen. This study did not generate new unique reagents.

### Experimental Model And Subject Details

### Human primed pluripotent stem cell culture

hESC cell line H9 (female) or H1 (male) were maintained on Matrigel (Corning) coated dishes in ncTarget hiPSC/hESC medium (Nuwacell) at 37℃ and passaged using Accutase to singlets or Dispase to colonies (Stem Cell Technologies). For maintenance, cells were passaged every 5 days.

For the pluripotency exit assay, hESCs were digested using Accutase (Stem Cell Technologies) and pipetting several times to harvest single cells, 80,000 hESCs were seeded into each well of a Matrigel (Corning) coated 12-well plate in ncTarget hESC medium (Nuwacell) supplemented with 10 μM Y-27632 (Selleck). 24 hours after seeding, hESC medium was replaced with TGFβ pathway inhibition medium (ncTarget medium with 1 mM A8301 (Tocris). Cells were incubated for 72 hours under TGFβi treatment. The medium was then replaced with ncTarget medium (Nuwacell) and incubated for 24 hours before use.

For embryoid bodies (EBs) formation assay, 6-well plates were coated using an anti-adherence rinsing solution (Stem Cell Technologies) for 30min at 37℃ to make the low-attachment plates. hESCs grown on Matrigel (Stem Cell Technologies) coated plate were digested using 1x collagenase IV (Thermo Fisher Scientific) for 45-60min at 37℃, 8ml basic DMEM medium (TransGen Biotech) were added to terminate digestion and pipetting softly for several times. The dissociated colonies were transferred to 15ml Falcon tubes for sedimentation and resuspended by EB medium, the EB pellets were transferred to a pre-treated low-attachment plate for culture (for one well of 6-well plate, two wells hESCs of 6-well plate is needed). Cell aggregates in suspension were gently cultured in EB medium and the plate was changed every other day. Differentiating EBs were harvested for RNA extraction on day 7. EB medium is composed of DMEM/F12 medium (Invitrogen), 20% Embryonic Stem Cell FBS, 1X Glutamax (Thermo Fisher Scientific), 1X MEM non-essential amino acids (Thermo Fisher Scientific) and 1X 2-Mercaptoethanol (Thermo Fisher Scientific), Primocin (100 µg/ml) (invivogen) and Penicillin-Streptomycin (100 U/mL) (Thermo Fisher Scientific).

### Human mesoderm cell induction

Human mesoderm cells were derived from hESCs using the STEMdiff mesoderm induction medium (Stem Cell Technologies) following the manufacturer’s instructions. Briefly, hESCs were digested using Accutase and pipetting several times to harvest single-cells. 50 000 cells per cm^2^ were seeded into Matrigel coated plates (for one well of 6-well plate, 5*10^5^ cells/well is needed), cells were incubated in ncTarget hESC medium (Nuwacell) supplemented with 10 μM Y-27632 (Selleck) at 37℃ for 24h, then medium were replaced with STEMdiff mesoderm induction medium for 5 days, mesoderm induction medium was changed every day, mesoderm cells were induced at about 72∼120 hours.

### 293T cell culture

293T (human, female) were obtained from ATCC (Cat# CRL3216) and maintained at 37℃ in DMEM medium (TransGen Biotech) supplemented with 10% FBS (Excell Bio), 1X Glutamax (Thermo Fisher Scientific), 100 mM MEM non-essential amino acids (Thermo Fisher Scientific), Primocin (100 µg/ml) (invivogen) Penicillin-Streptomycin (100 U/mL) (Thermo Fisher Scientific).

### Generation of mammal cells with inducible LSM14A-GFP constructs

Human LSM14A-GFP (Ensembl Transcript ID: ENST00000544216.8) were cloned into FUW-tetO-loxP vector (was a gift from Rudolf Jaenisch lab, Addgene #60849) via Gibson assembly, for preparation of lentiviruses, 293T cells in 6-well plates were transfected at 80% confluency with the lentiviral vector 3200 ng FUW-tetO-loxP containing LSM14A-GFP, the lentiviral packaging plasmids 800 ng pMD2.G (was a gift from Didier Trono, Addgene #12259) and 2400 ng psPAX2 (was a gift from Didier Trono, Addgene #12260), with lipofectamine 2000 for 24h. FUW-M2rtTA (a gift from Rudolf Jaenisch lab, Addgene #20342) lentivirus containing rtTA were prepared in the same way. 48 hours after transfection, the cell medium containing lentivirus was harvested and filtered through a 0.45mm filter followed by concentration using Lenti-Concentrator (Genomeditech). 5*10^5^ cells were infected with the two lenti-virus and cultured in medium (with 10μM Y-27632 for hESC) for 120h. Single cells were sorted into 96-well plate for picking individual colonies. Obtained clones were genotyped by PCR and sequenced and GFP intensity and homogeneity were verified using flow cytometry and immunofluorescence image. Candidate clones under doxycycline (1μg/ml for hESC and 100ng/ml for 293T) (Selleck) induction with typical p-body characters were used for P-body purification.

## METHOD DETAILS

### P-body purification by fluorescent associate particle sorting

P-body purification was performed as described in Hubstenberger et al., 2017^19^ with little modification. Briefly, 293T, hESCs and hESC-derived mesodermal cells carrying tetO-LSM14A-GFP were under doxycycline (1μg/ml for hESCs and 100ng/ml for 293T) (Selleck) induction for 4 days and grown to 90% confluency in ten 15 cm dishes. For 293T, cells were washed twice in pre-chilled PBS and scrapped in 2 mL PBS, for hESCs and mesodermal cells, cells were washed twice in PBS and digested by trypsin-EDTA (0.05%) (Thermo Fisher Scientific), then washed by pre-chilled PBS. The following steps are all performed on ice or at 4℃, cell pellets were suspended in lysis buffer (50 mM Tris, pH 7.4, 1 mM EDTA, 150 mM NaCl, 0,2% Triton X-100) with 65 U/mL RNaseOut ribonuclease inhibitor (Roche Diagnostics) and EDTA-free protease inhibitor cocktail (Roche Diagnostics). Extracts were passed 20 times through a 27G syringe needle (BD Biosciences), followed by a total incubation time about 30min. Lysates were spun for 5 min at 200g to deplete nuclei, followed by supplemented with 10 mM MgSO4 and 1 mM CaCl2 and treated with 4 U/mL of RQ1 DNase (Promega) for 30 min at RT and spun down for 10 min at 10000g, pellets were resuspended into 40 μL of lysis buffer with 80 Units of RNaseOut ribonuclease inhibitor (Roche Diagnostics). This pre-sorted fraction was named cytoplasm, from which P-bodies were sorted by Influx Cell Sorter (BD Biosciences) with a nozzle size of 70 mm. Particles were detected according to the Forward-scattered light (FSC) and the GFP fluorescence using the 488 nm excitation laser and the 526/52 band pass filter. The P-body sorting size was defined by 3µm Sphero Rainbow Calibration Particles (BD Biosciences) and the GFP intensity was defined by tetO-NES-GFP cells, and additional channels PerCP (laser: 488, filter: 670/30) and PE (laser: 561, filter: 585/29) were introduced to remove the non-specific fluorescent of debris. Purity mode (1-2 envelope) was applied for sorting. Cytoplasm was diluted using lysis buffer before sorting and the dilution rate is based on the flow sorting events rate. About 5*10^6^ P-bodies were collected for 4 hours of sorting. The collected GFP+ fraction was spun for 10min at 10000g to pellet P-bodies, P-bodies and the corresponding cytoplasm were lysate in TRIzol LS (Invitrogen) for the following RNA isolation and sequencing.

### RNA sequencing

Total RNA of P-bodies and corresponding cytoplasm or cellular samples were isolated using TRIzol LS (Invitrogen) following the manufacturer’s instructions. The mRNA fraction was enriched by pulldown of poly(A)-RNA using mRNA capture beads (Vazyme) and RNA sequencing libraries were generated using Universal V8 RNA-seq Library Prep Kit for Illumina (Vazyme) according to the manufacturer’s protocol followed by sequencing using Illumina NovaSeq-6000 platform with 150 bp paired-end reads (Novogene). Libraries’ qualities were assessed by qPCR and the Agilent 2100 Bioanalyzer instrument (Agilent).

### Western Blot

The sorted P-bodies and corresponding cytoplasm were spun down at 10000g for 10min and lysed in 50μL RIPA buffer (50 mM Tris, 150 mM NaCl, 0.1% SDS, 0.5% sodium deoxycholate, 1% Triton X-100, pH 7.5) supplemented with 1x protease inhibitor cocktail and 1 mM PMSF, combined with protein loading buffer and boiled for 10 min. The resolved proteins were subjected to immunoblot analyses using primary antibody: Rabbit-anti-human-DDX6 (1:1000, Huabio, ET7107-06) and secondary antibody: Donkey-anti-rabbit-IgG-HRP (1:2000, Cell Signaling Technology, 7074s). Proteins were detected using the WesternBright ECL kit (Advansta), and visualized by Odyssey Fc Imaging System (LI-COR Biosciences).

### Immunofluorescence and fluorescence microscopy

Cells for immunofluorescence assay were grown on glass bottom cell culture dishes (NEST) and fixed in 4% PFA for 10 min, and then incubated with the primary antibody for 1 hour, followed by incubated with the fluorochrome-conjugated secondary antibody for 1 hour, mounted in ProLong gold antifade mountant with DAPI (Sigma-Aldrich). All steps were performed at room temperature and cells were rinsed with PBS for three times at the interval of each step. Confocal microscopy was performed on LSM 880 with Airyscan (ZEISS). Primary antibodies used in this study include: rabbit-anti-DDX6 (1:200, Huabio, ET7107-06), rabbit-anti-DCP1A (1:200, Huabio, ET7107- 26), mouse-anti-OCT4(1:200, BD Biosciences, 611203), rabbit-anti-NANOG (1:200, Cell Signaling Technology, 4903S), rabbit-anti-SOX2(1:200, Abcam, ab97959), rabbit-anti-BRACHYURY (1:200, Cell Signaling Technology, 81694S). Secondary antibodies were purchased from Abcam or Invitrogen. Images were processed using ImageJ.

### RNAscope

For In situ RNA hybridization, the probe targeting SPEN (Advanced Cell Diagnostics, 419971) was designed and validated. RNAscope Multiplex Fluorescent Reagent Kit v2 (Advanced Cell Diagnostics, #323100) was used for all FISH experiments according to the manufacturer’s protocol. Briefly, cell samples were fixed in 10% formalin solution neutral buffered (RNase Free) at room temperature for 30 mins, treated with RNAscope hydrogen peroxide for 10 min, treated with RNAscope protease III for 15 min at 40℃. The cellular samples were incubated with mRNA probes for 2 hours at 40 °C and specific signals were amplified with multiplexed implication buffer and detected with TSA Plus Fluorescent Kit (Advanced Cell Diagnostics, 322809). Samples are rinsed with 1x wash buffer for three times at the interval of each step. The confocal microscopy and the imaging process are the same as described above.

### Vectors and cloning

For the DDX6-GFP knock-in construct, ∼1000bp upstream near the genome of DDX6 stop codon fused with GFP were ligated into PL552 vector with puromycin resistance (was a gift from Su-Chun Zhang, Addgene #68407) through EcorI restriction enzyme recognition sites via Gibson assembly. ∼1000bp downstream near the genome of DDX6 stop codon sequence was ligated into the construct through BamHI sites via Gibson assembly. gRNA sequences were designed using the CHOPCHOP gene-editing tool (http://chopchop.cbu.uib.no/) to be located close to the stop codon (to enable C-terminal fusions in PL552-DDX6-P2A-GFP with homologous arms), annealed oligos with BbsI sites encoding corresponding gRNAs were ligated into PX459 (was a gift from Feng Zhang, Addgene #62988). The PL552-DDX6-P2A-GFP and PX459-sgRNA constructs were transfected into the hESCs with P3 Primary Cell 4D Nucleofector X Kit S (Lonza) on 4D-nucloefector (program CA137, Lonza) following the manufacture’s protocol. 8*10^5^ hESCs and 4μg of each plasmid were used for nucleofection. Nucleofected cells were then seeded in a single well of a 6-well plate in ncTarget hESC medium supplemented with 10μM Y-27632 (Selleck) for 72h. Puromycin selection (1 μg/ml) was applied post-nucleofection for another 72h, 10000 cells were seed to a Matrigel-coated 10 cm dish with Y-27632 (10 μM) for 48h after drug selection until stable colonies appeared, and colonies were picked and passaged at least two times. The obtained clones were again genotyped by PCR and sequenced. GFP intensity and homogeneity were verified using flow cytometry and IF.

For KRAB-BFP stable construct, pC13N-dCas9-BFP-KRAB (Addgene #127968), CLYBL-L, CLYBL-R (all were gifts from Ruilin Tian lab) constructs were transfected into the hESCs with P3 Primary Cell 4D Nucleofector X Kit S (Lonza) on 4D-nucloefector (program CA137, Lonza) followed the manufacturer’s protocol. 8*10^5^ hESCs and 4μg of each plasmid were used for nucleofection. Nucleofected cells were then seeded in a single well of a 6-well plate in ncTarget hESC medium supplemented with 10μM Y-27632 (Selleck) for 96-120h. BFP positive cells were sorted by Influx cell sorter (BD Biosciences), the sorted cells were amplified and repeated for sorting until the positive rate reached 100% BFP positive. The KRAB-BFP stable expression clones were again genotyped by PCR and sequenced, and BFP intensity and homogeneity were verified using flow cytometry and IF. Of note, KRAB-BFP is inserted into the genome of CLYBL stop codon and does not impact hESC function.

For CRISPRi knock-down construct, gRNA sequences were obtained from a previous study^55^, two annealed oligos with BbsI sites encoding corresponding gRNAs were ligated into modified plenti-H1-U6-mCherry (was a gift from Chaochen Wang lab) via Gibson assembly. 293T cells in 6-well plates were transfected at 80% confluency with 3200ng lentiviral vector plenti-H1-U6-mcherry with dual sgRNA, 800ng lentiviral packaging plasmids pMD2.G (was a gift from Didier Trono, Addgene #12259) and 2400ng psPAX2 (was a gift from Didier Trono, Addgene #12260), with Lipofectamine 2000 for 24h. FUW-M2rtTA (was a gift from Rudolf Jaenisch lab, Addgene #20342) lentivirus containing rtTA were prepared in the same way. About 48 hours after transfection, the cell medium containing lentivirus was harvested and filtered through a 0.45-mm filter followed by concentrated using Lenti-Concentrator (Genomeditech). 5*10^5^ hESC with KRAB-BFP stable expression were infected and cultured in medium with 10μM Y-27632 for 120h, and mCherry positive cells were sorted by Influx cell sorter (BD Biosciences). The obtained clones were again genotyped by qRT-PCR. mCherry intensity and homogeneity were then verified using flow cytometry and IF.

### qRT-PCR analyses

Total RNA for qRT-PCR was isolated using RNA isolator (Vazyme) following the manufacturer’s instructions, and quantified using Nanodrop One (Thermo Fisher Scientific). cDNA was synthesized with PrimeScript RT reagent Kit with gDNA Eraser (TaKaRa). qRT-PCR reactions were set up in at least triplicate with the AceQ qPCR SYBR Green Master Mix (Vazyme). Reactions were run on the LightCycler 480 System (Roche Diagnostics). Primers are available upon request from authors.

### Flow cytometry analysis

Cells were analyzed with Cytek Aurora Flow Cytometry (Cytek Biosciences) using SpectroFlo. 7-AAD or DAPI was used to indicate the viability of the cells. Antibodies used in this study include: EpCAM conjugated with AF488 (1:200, Biolegend, B275293). Analysis and visualization of the flow cytometry data was performed by FlowJo (v10.0.7).

### Cycloheximide test

293T cells grown on glass bottom cell culture dishes (NEST) at 80% confluency were treated with 100 µg/mL of cycloheximide for 4 hours, rinsed with PBS twice, and replaced with culture medium. Microscopy was performed on the inverted fluorescence microscope NIB900 (NOVEL).

### Fluorescence recovery after photobleaching

FRAP experiments were performed when cells were grown on glass bottom cell culture dishes (NEST) at 80% confluency, using the LSM 880 with Airyscan (ZEISS). Experiments were performed at 37˚C under 5% CO_2_ using a live-cell chamber system. Cells were imaged in the GFP channel for the detection of LSM14A (P-bodies), and at least nine z-slices were taken every 400 nm. The target P-body was bleached for 500ms using the 488 nm laser. Six pre-bleach images were acquired. Images were taken every 2s. Images were analyzed using ImageJ.

## QUANTIFICATION AND STATISTICAL ANALYSIS

### Statistical analysis

Quantitative data are presented as means ± SEM. Statistical analyses were performed using Prism 9 (GraphPad) or R-project. Details for statistical analyses, including replicate numbers, are included in the figure legends.

### RNA-seq analysis

Qualities of raw RNA-seq reads were assessed using fastQC (RRID:SCR_014583). Extra adaptors and low-quality reads were removed using Cutadapt^59^ and Trim Galore(RRID:SCR_011847). clean reads were mapped to the GRCh38 Ensembl genome with Homo_sapiens.GRCh38.110.gtf annotations (https://ftp.ensembl.org/pub/release-110/gtf/homo_sapiens/) using STAR ^60^ followed by Bam converted, sorted and indexed using SAMtools^61^, and were counted using featureCounts^62^ by comprehensive gene annotation (human v44, GENCODE) for mRNA, long non-coding RNA gene annotation (human v44, GENCODE) for lncRNA and repeat sequence annotation for repeat sequences (https://repeatbrowser.ucsc.edu/).

Differential expression analysis was performed using DESeq2^63^ after normalization, transcripts with TPM (CPM for repeat sequences) <1 in all samples were removed. Differentially expressed genes were determined based on the criteria of > 1 log2 fold change for comparison in expression value and adjust p-value < 0.05. For further analysis, DEGs that exclusively had ENSG IDs (primarily representing novel transcripts) were excluded. PCA, chord, heatmap and scatter plots were visualized by ggplot2^69^, circlize^70^ and ComplexHeatmap (RRID:SCR_017270) packages in R-project.

### GC content, length, and m6A modification analysis

For GC content and length analysis, all GC content and length data were acquired using the Ensembl biomaRt package^68^ in R-project, and isoforms less than 500bp were omitted. For m6A modification analysis, m6A modified sites data were acquired from a previous report^32^, numbers and sites of m6A modification in P-body enriched or depleted genes were calculated in R-project.

### mRNA functional enrichment and Protein-protein interaction analysis

Functional enrichment, enriched clusters and PPIs analysis were generated using Metascape online service (https://metascape.org). PPIs were verified in the STRING database (https://string-db.org). The enriched clusters and PPI networks were imported into Cytoscape (www.cytoscape.org).

### Protein localization and key stem cell functional enrichment analysis

Although P-body transcripts are mainly translationally repressed mRNAs associated with regulatory processes ^10^, the composition of the transcripts and the potential regulated function are still largely unclear. Considering that mRNA-encoded proteins may be involved in specific cellular functions, the subcellular localization of proteins, as well as GO terms or KEGG signaling pathways associated with embryo development, its association with enriched or depleted in P-body may potentially reflect P-body regulatory functions.

For protein localization analysis, we used subcellular protein localization data from a previous study^33^. Calculated the enrichment score for each sample in each location according to the normalized formula: background ratio = enriched proteins of specific location/all proteins, gene ratio = genes enriched in specific protein location term/all input genes, enrichment score = gene ratio/background ratio. We scaled the enrichment score to localization to better represent differences between samples. Stem cell associated terms analysis was carried out using clusterProfiler package^65^ in R-project, and visualized by scaled enrichment score to terms.

### lncRNA interactions analysis

Due to the incomplete annotation of lncRNA functions, lncRNA-associated potential regulators and targets including RNA and proteins were evaluated using an integrated tool lncSEA^67^ (https://bio.liclab.net/LncSEAv2/index.php). ENCORI, RNAInter4.0, and NPinter4.0 databases were used to determine the lncRNA-protein interaction. RISE, ENCORI, RNAInter4.0, NPinter4.0, and LncBase2.0 databases were used to determine the lncRNA-RNA interaction, and KnockTF database was used to determine the transcription factors perturbation to elucidate the potential TF-related functions.

### Repeat sequence analysis

Repeat sequences including rRNA and transposable elements (TEs) were analyzed in this study, CPM (count per million) normalization was used to facilitate comparisons of TE classes/families. Differential expression analysis was performed using DESeq2^63^ as described above.

## DATA AND CODE AVAILABILITY

All relevant data are included in the paper or Supplementary Information files and are available from the authors. The RNA-seq data generated in this paper is available at GEO: GSE246537. The pipeline for the project can be found on Zenodo (10.5281/zenodo.10050771).

## Notes

### Competing Interest Statement

The authors have declared no competing interest.

